# The chromatin landscape of Th17 cells reveals mechanisms of diversification of regulatory and pro-inflammatory states

**DOI:** 10.1101/2022.02.26.482041

**Authors:** Pratiksha I. Thakore, Alexandra Schnell, Maryann Zhao, Linglin Huang, Yu Hou, Elena Christian, Sarah Zaghouani, Chao Wang, Vasundhara Singh, Sai Ma, Venkat Sankar, Samuele Notarbartolo, Jason D. Buenrostro, Federica Sallusto, Nikolaos A. Patsopoulos, Orit Rozenblatt-Rosen, Vijay K. Kuchroo, Aviv Regev

## Abstract

Th17 cells are a heterogenous cell population consisting of non-pathogenic Th17 cells (npTh17) that contribute to tissue homeostasis and pathogenic Th17 cells (pTh17) that are potent mediators of tissue inflammation. To reveal regulatory mechanisms underlying Th17 heterogeneity, we performed combined ATAC-seq and RNA-seq and discovered substantial differences in the chromatin landscape of npTh17 and pTh17 cells both *in vitro* and *in vivo*. Compared to other CD4^+^ T cell subsets, npTh17 cells share accessible chromatin programs with T_regs_, and pTh17 cells have an intermediate profile spanning features of npTh17 cells and Th1 cells. Integrating single-cell ATAC-seq and single-cell RNA-seq, we inferred self-reinforcing and mutually exclusive regulatory networks controlling the different cell states and predicted transcription factors (TFs) shaping the chromatin landscape of Th17 cell pathogenicity. We validated one novel TF, BACH2, which promotes immunomodulatory npTh17 programs and restrains pro-inflammatory Th1-like programs in Th17 cells and showed genetic evidence for protective variants in the human BACH2 locus associated with multiple sclerosis. Our work uncovered mechanisms that regulate Th17 heterogeneity, revealed shared regulatory programs with other CD4^+^ T cell subsets, and identified novel drivers of Th17 pathogenicity as potential targets to mitigate autoimmunity.

## Introduction

Balancing proinflammatory and regulatory CD4^+^ T helper subsets is essential for an effective immune response towards pathogens, while avoiding uncontrolled inflammation and autoimmunity. Among those, IL-17-producing CD4^+^ T cells (Th17 cells) play a key role in the host defense against extracellular pathogens and contribute to mucosal barrier homeostasis ^1, 2^. However, Th17 cells are also important pathogenic drivers of multiple autoimmune diseases, including multiple sclerosis (MS), psoriasis, and Sjogren’s syndrome ^3–5^.

Mirroring this functional diversity, *in vitro* polarization of naïve CD4^+^ T helper cells with IL-6 and TGF-ß yields Th17 cells that instigate little or no tissue inflammation or autoimmunity (non-pathogenic Th17 cells (npTh17)), while IL-6, IL-1ß and IL-23 jointly generate Th17 cells highly potent in transferring disease (pathogenic Th17 cells (pTh17)) ^6, 7^. Previous work, focused on the expression programs that distinguish Th17 cell populations, revealed distinct gene signatures active in npTh17 cells *vs*. pTh17 cells ^6, 8^. While npTh17 cells express the immunoregulatory genes *Il10*, *Il9*, *Maf*, and *Ahr*, pTh17 cells express proinflammatory genes *Csf2*, *Ifng*, *Tbx21, Il23r* and *Gzmb*. Similar subsets of Th17 cells have also been identified in humans, where Th17 cells express either IL-10 or IFNγ, depending on whether they are specific for *Staphylococcus aureus* or *Candida albicans*, respectively ^9^.

Th17 cells demonstrate additional plasticity ^4, 10, 11^, with a capacity to acquire either beneficial functions associated with other Th cells or pathogenic pro-inflammatory states, often characterized by the expression of the Th1-cytokine IFNγ ^12–16^. For example, in the intestines Th17 cells trans-differentiate into IL-10-producing T regulatory 1-like cells (Tr1-like cells) ^17^. Conversely, in models of autoimmunity, such as experimental autoimmune encephalomyelitis (EAE), T cells that initially express IL-17 (ex-Th17 cells) upregulate the expression of IFNγ and GM-CSF ^12, 16^, and the transition of Th17 cells to Th1-like cells is required for their ability to induce colitis in a transfer model ^13, 18^. In human autoimmune diseases, T cells co-expressing IL-17A and IFNγ are correlated with pathogenicity ^10, 19–23^.

The extent to which npTh17 and pTh17 populations represent transient cell states or stable cell fates remains elusive, as is the relationship of pTh17 cells expressing Th1-lineage genes, such as *Tbet* and *Ifng*, to *bona fide* Th1 cells. In particular, it remains unclear whether pTh17 and npTh17 cells are distinct, stable cell fates or are functional states, wherein Th17 cells toggle between the two states. In addition, the master regulators of different Th17 programs and their plastic transitions have yet to be comprehensively deciphered. Understanding these molecular switches can help design therapeutic strategies that specifically target pathogenic, disease-driving Th17 populations yet retain non-pathogenic homeostatic Th17 cell function in barrier integrity and prevention of microbial invasion at mucosal surfaces.

Accessible chromatin distinguishes active regulatory elements that drive gene transcription and define cell state, and chromatin accessibility profiles can stably distinguish cell lineages and foreshadow RNA expression changes during differentiation ^24–27^. Common genetic variants in cell-type specific distal regulatory elements in immune cells have been associated to human autoimmune diseases, such as MS and inflammatory bowel disease (IBD) ^28^. Thus, we hypothesized that comparing the chromatin states of npTh17 and pTh17 cells can help determine if Th17 functional plasticity is marked by distinct epigenomic features and identify regulators of the pTh17 cell fate.

Here, we characterized the chromatin accessibility and associated expression profiles of *in vitro* and *in vivo*-derived npTh17 and pTh17 cells with ATAC-seq and RNA-seq in cell populations and single cells. We found highly distinct chromatin landscapes in npTh17 and pTh17 cells, such that npTh17 and pTh17 cells each also shared substantial accessible chromatin programs with T_reg_ and Th1 cells, respectively. Integrating single-cell ATAC-seq (scATAC-seq) and single cell RNA-seq (scRNA-seq) allowed us to map chromatin signatures of Th17 pathogenicity to effector genes and to predict multiple novel TF regulators of the pathogenic Th17 program. Among these, we highlight BACH2, which we validated as a novel regulator of the Th17 chromatin landscape and Th17 pathogenicity *in vitro* and *in vivo.* Our analysis provides insights into the regulatory programs of Th17 cells, the relationship of Th17 cells to other CD4^+^ T cell subsets and identifies novel drivers of the Th17 pathogenic program.

## Results

### Distinct chromatin landscapes for pathogenic and non-pathogenic Th17 cells

We hypothesized that the distinct pro-inflammatory expression programs of pTh17 *vs*. npTh17 cells ^6, 8^ would be reflected in their respective epigenetic landscapes. To test this hypothesis, we performed ATAC-seq of *in vitro* differentiated npTh17 and pTh17 cells at 24h, 48h, and 72h after stimulation of naïve Th cells with TGF-β1 and IL-6 or IL-1β, IL-6 and IL-23, respectively (**Fig. 1A**). To assess the relation of IL-17A expression to the chromatin landscape, we also compared Th17 cells selected for IL-17A-GFP expression to all CD4^+^ cells, using an *Il17a*^GFP^ reporter mouse (**Fig. S1A**).

**Fig. 1:**
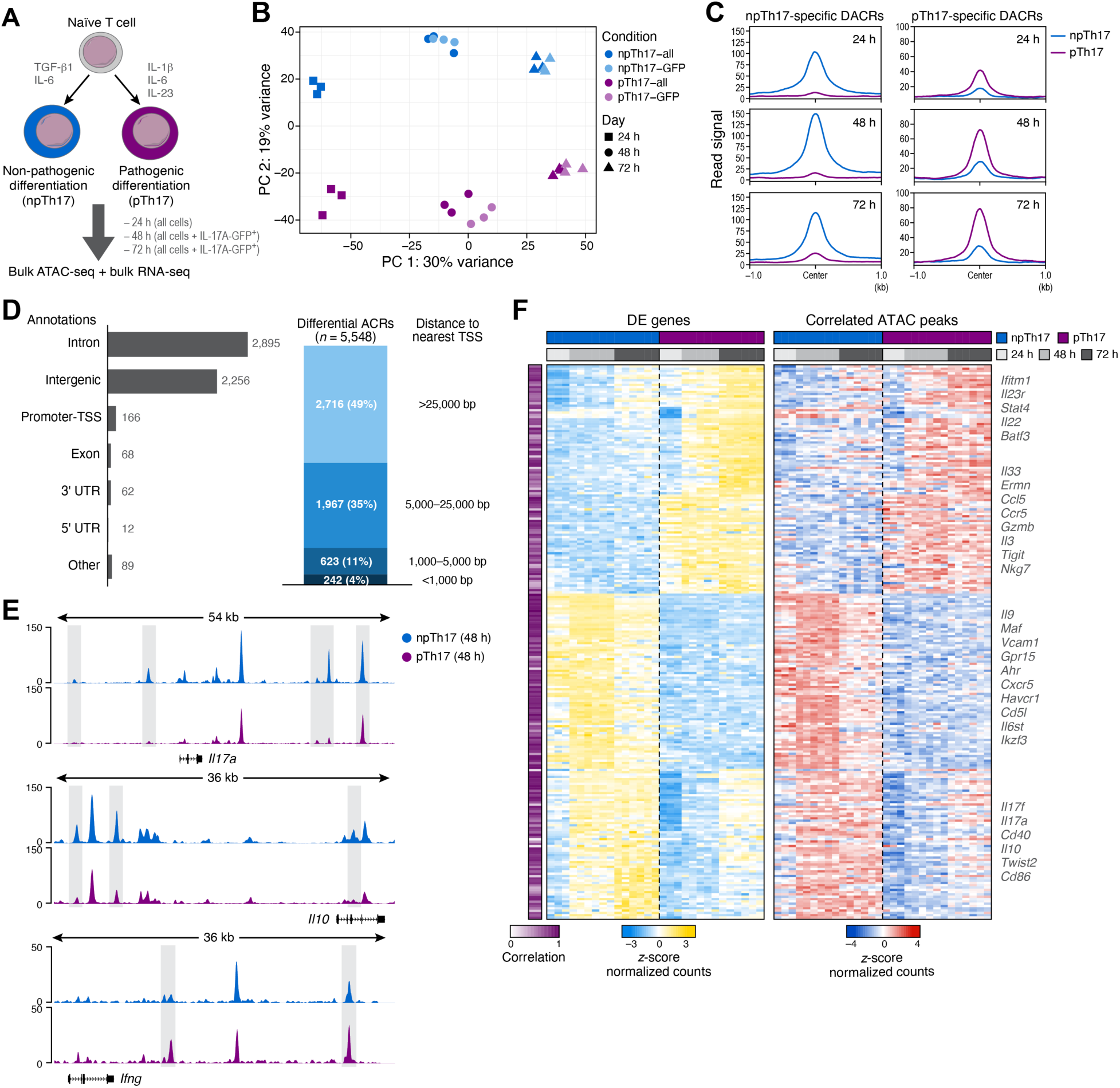
Distinct chromatin landscapes in non-pathogenic and pathogenic Th17 cells. (**A**) Experimental setup of *in vitro* system. Naïve CD4^+^ CD25^-^ CD44^-^ CD62L^+^ T cells (grey, top) were *in vitro* differentiated to npTh17 (blue, left) or pTh17 (purple, right) cell populations. (**B**,**C**) Distinct chromatin accessibility profiles for pTh17 and npTh17 cell populations. (**B**) First (*x* axis, PC1) and second (*y* axis, PC2) principal components of a PCA of ATAC-seq profiles (dots) of *in vitro* differentiated cell from all (dark color) or GFP^+^ (light color) cells sorted from npTh17 (blue) and pTh17 (purple) populations at 24h (square), 48h (circle), and 72h (triangle) after polarization of naïve CD4^+^ T cells from *Il17a*^GFP^ reporter mice. (**C**) Global chromatin accessibility (*y* axis, normalized reads) of npTh17 (blue) and pTh17 (purple) cell populations in chromatin regions that are more accessible in npTh17 (left) or in pTh17 (right) populations. (D) Differentially accessible chromatin regions (DACRs) are enriched in intronic and intergenic regions. Number of DACRs between *in vitro* differentiated npTh17 and pTh17 cells 48h after polarization (left, *x* axis) at different annotation locations (left, *y* axis) and their proportion (right, *y* axis) at different distances to nearest TSS (right). (E) Differentially accessible chromatin regions between npTh17 and pTh17 populations at key Th17 cell effector loci. Enrichment of ATAC-seq signal over background (*y* axis, log fold) for npTh17 (blue) and pTh17 (purple) cell populations profiled at 48h along the *Il17a* (top, *x* axis, chr1:20,710,909-20,765,000), *Il10* (middle, *x* axis, chr1:130,989,883-131,026,008), and *Ifng* (bottom, *x* axis, chr10:118,439,591-118,475,730) loci. (F) Agreement in gene expression and chromatin accessibility profiles. Expression levels (left, z-score of normalized counts, color bar) and ATAC counts at the peak 100kb upstream or downstream of gene body (right, z-score of normalized counts, color bar) for differentially expressed genes (rows) with the highest correlation of accessibility signal with gene expression. Left purple column: Pearson’s correlation coefficient between gene expression and ATAC-seq.

Principal component analysis (PCA) revealed substantial differences in the chromatin landscape between npTh17 and pTh17 cells, consistently across time points (**Fig. 1B,C**). While principal component 1 (PC1) reflected changes over time consistently in both npTh17 and pTh17 cells, PC2 separated npTh17 from pTh17 cells at each time point (**Fig. 1B**), showing that the two types of Th17 cells diverge into two cell fates at least as early as 24h after treatment with polarizing cytokines. The chromatin landscape of IL-17A-GFP Th17 cells largely resembled Th17 cells that were not selected for IL-17A expression (**Fig. 1B**), with only a few differences (65 differentially accessible chromatin regions (DACRs), FDR <0.05). Hence, for all subsequent comparisons, we used Th17 cells that were treated with polarizing cytokines, but not sorted for IL-17A-GFP. Overall, 5,685, 7,039, and 3,958 peaks were differentially accessible between npTh17 and pTh17 cells at 24h, 48h, and 72h, respectively (11,346 unique peaks total, 1,090 shared across all three time points).

We focused on DACRs at 48h post-polarization, where we observed the greatest number (7,039) of differentially accessible peaks between pTh17 and npTh17 cells (**Fig. 1C**). The vast majority of DACRs were located in non-coding regions (intronic and intergenic), distally (>25K bp) from the nearest transcription start site (TSS) (**Fig. 1D**). Of the 7,039 DACRs detected at 48h, 4,645 were more accessible in npTh17 cells and 2,394 were more accessible in pTh17 cells. These includes DACRs at sites proximal to Th17 effector genes, including *Il17a* and *Il17f* (higher in npTh17), genes associated with regulatory Th17 programs (*e.g.*, *Il10;* higher in npTh17), and genes associated with pro-inflammatory Th17 gene programs (*e.g.*, *Ifng and Csf2;* higher in pTh17) (**Fig. 1E**, shaded boxes). Overall, these results show that polarized pTh17 cells represent a distinct cell state from npTh17 cells, characterized by unique chromatin accessibility profile, including in key loci.

### Chromatin landscape changes correspond to expression differences between npTh17 and pTh17 cells

There was a strong positive correlation between the differences between npTh17 and pTh17 cells in the chromatin landscape (as reflected by DACRs) and the differences in gene expression programs (by differentially expressed genes). Specifically, to relate differences in chromatin accessibility with changes in Th17 gene programs, we collected matched RNA-seq from *in vitro* polarized Th17 cell populations (**Fig. S1B,C**), and searched a 100kb window upstream and downstream of the gene body of the top 200 genes differentially expressed between npTh17 and pTh17 cells at each time point (FDR <0.05, ranked by fold-change, 416 genes) for the chromatin peak whose ATAC-seq signal correlated most with each gene’s expression across conditions (**Fig. 1F** and **Table S1**). Of the 416 genes tested overall, 240 had a nearby DACR with a significant positive correlation (average Pearson’s *r* = 0.70, FDR <0.05).

The npTh17-specific genes with correlated chromatin changes included immunoregulatory genes previously associated with the npTh17 phenotype (*Il9*, *Maf*, *Ahr*, *Cd5l*, *Il10*) ^6, 8, 29^, as well as genes not yet associated with npTh17 cells (*Gpr15*, *Havcr1*, *Twist2*) but with previously defined regulatory functions in other immune cell types ^30–32^. The pTh17-specific genes included established proinflammatory genes (*Il23r*, *Stat4*, *Batf3*, *Gzmb*, *Nkg7*) ^6, 8^ and novel genes (*Il33*, *Ermn*) not previously implicated in the pTh17 phenotype or function. Interestingly, several key loci with changes in chromatin accessibility (*e.g. Ifng* and *Csf2*) did not have corresponding changes in expression, suggesting that genes in these loci are poised for transcription by a permissive chromatin context. Overall, these results indicate that pathogenic and non-pathogenic Th17 cells have distinct epigenetic regulatory mechanisms that are associated with pro-inflammatory and regulatory gene programs.

### Distinct chromatin landscapes of Th17 cells from the dLN and CNS during EAE are partly mirrored by npTh17 and pTh17 cells *in vitro*

To relate the chromatin landscapes of npTh17 and pTh17 cells to *in vivo* states, we next collected ATAC-seq profiles of Th17 cells from EAE, a mouse model for MS and an *in vivo* model of Th17-mediated autoimmunity. Previous scRNA-seq showed similarities between the cell states of *in vitro* differentiated npTh17 cells *vs*. pTh17 cells and those of draining lymph node (dLN)-derived Th17 cells *vs*. central nervous system (CNS)-derived Th17 cells in EAE mice ^8^. Hence, we induced EAE in *Il17a*^GFP^ reporter mice, sorted viable IL-17A-GFP^+^ CD4^+^ T cells from the dLN and the CNS at peak of disease, and profiled them by bulk ATAC-seq and RNA-seq (**Fig. 2A-C, S2A-C** and **Table S2, S3**).

**Fig. 2:**
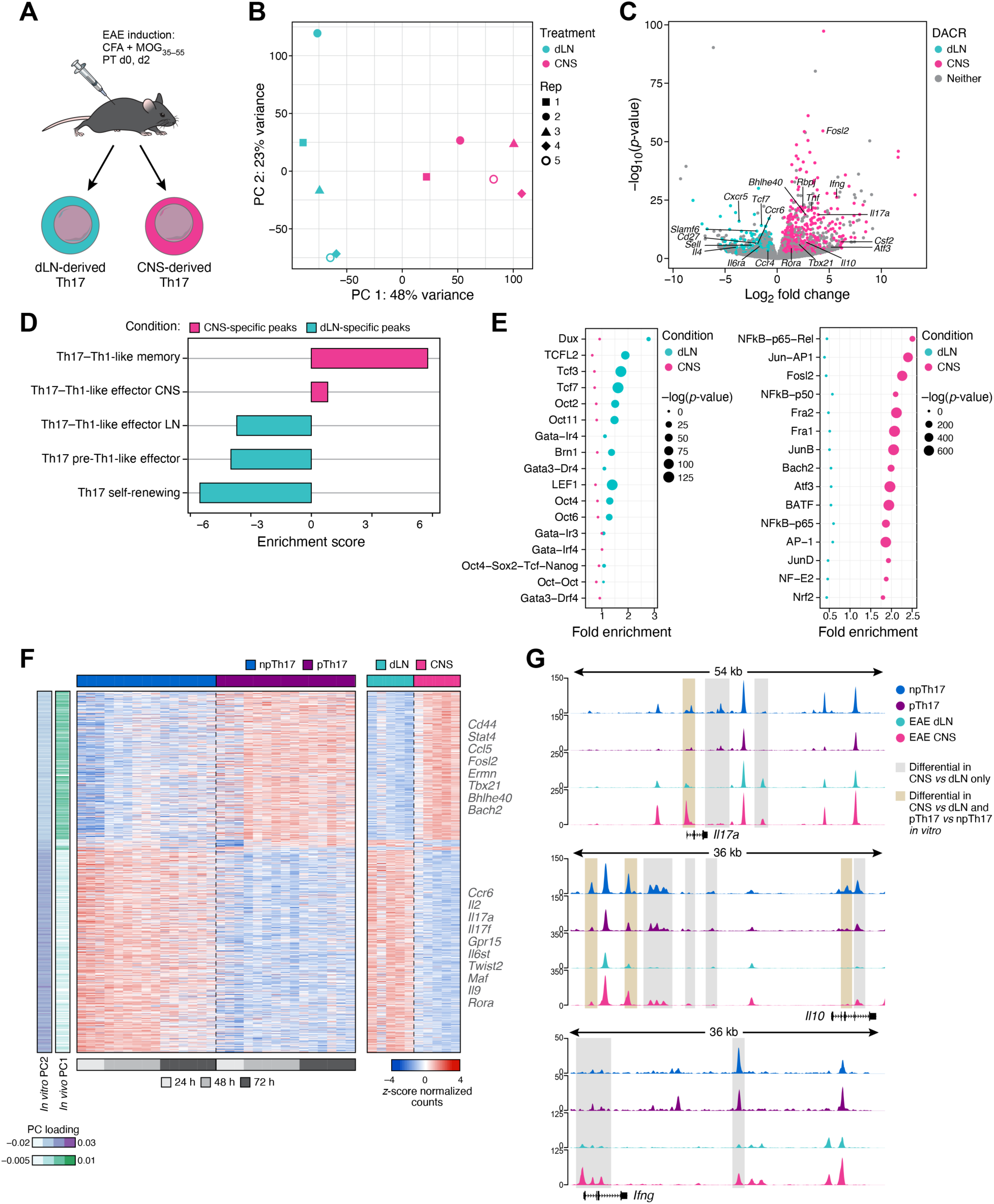
Distinct chromatin accessibility profile of CNS-infiltrating Th17 cells during EAE. (**A**) Experimental setup of *in vivo* system. Active EAE was induced in *Il17a*^GFP^ reporter mice and Th17 cells were harvested from the dLN (teal, left) and CNS (pink, right) at peak of disease for bulk ATAC-seq and bulk RNA-seq. (**B**) Distinct chromatin accessibility profiles of CNS-infiltrating Th17 cells. First (*x* axis, PC1) and second (*y* axis, PC2) principal components of a PCA of ATAC-seq profiles (dots) of dLN-derived (teal) and CNS-derived (pink) IL17A-GFP^+^ cells. dLN and CNS samples are matched for each biological replicate (shapes, n=5). (**C**) Differentially expressed genes in CNS-infiltrating Th17 cells are associated with differentially accessible chromatin peaks. Differentially expression (*x* axis, log_2_(fold change)) and significance (*y* axis, -log_10_(p-value)) between CNS-derived and dLN-derived Th17 cells profiled by bulk RNA-seq. Differentially expressed genes with a corresponding differentially accessible chromatin peak (FDR <0.05 for CNS *vs*. dLN) are highlighted (teal for dLN, pink for CNS). (**D**) Chromatin accessibility changes in CNS-derived Th17 cells are enriched for pathogenic Th17 signatures ^8^. Enrichment score (*x* axis, normalized enrichment score) of genes associated with the dLN-specific (teal) and CNS-specific (pink) ATAC peaks for different Th17 and Th1 gene signatures (*y* axis) (**Table S4**). (**E**) TF motifs differentially enriched in dLN or CNS-derived Th17 cells. Significance (-log10(p-value), dot size) of fold enrichment (*x* axis) over background, of TF motifs (*y* axis) rank ordered by fold enrichment (*x* axis) in differential ATAC-seq peaks in Th17 cells from the dLN (left, teal) or CNS (right, pink). (**F,G**) Shared chromatin accessibility features between *in vitro*- and *in vivo*- derived Th17 cells. (F) Normalized counts (z-score, color bar) of DACRs (rows) shared between npTh17 and dLN-derived Th17 cells (bottom rows) or between pTh17 and CNS-derived Th17 cells (top rows). Selected genes associated with the ATAC-seq peaks are labeled on right. Color bars (left): PC loading from PCAs of ATAC-Seq of *in vitro* cells (PC2 of Fig. 1B; purple, left) and of *in vivo* cells (PC1 of Fig. 2B; cyan, right). (**G**) Enrichment of ATAC-seq signal over background (*y* axis, log fold) for npTh17 (blue, 72h), pTh17 (purple, 72h), dLN-derived (teal), and CNS-derived (pink) cell populations at the *Il17a* (top, chr1:20,710,909-20,765,000), *Il10* (middle, chr1:130,989,883-131,026,008), and *Ifng* (bottom, chr10:118,439,591-118,475,730) loci.

There were substantial differences in the chromatin landscape of Th17 cells from the dLN and CNS (24,694 DACRs of 137,282 total peaks assayed, FDR <0.05, **Fig. 2B,C** and **S2C**), with 13,076 and 11,618 DACRs specific to dLN and CNS Th17 cells, respectively. dLN-specific peaks were proximal to genes associated with the non-pathogenic Th17 phenotype (*Cxcr5*, *Sell*, *Tcf7*, *Il6ra, Maf*) ^6, 8^, whereas CNS-specific peaks were close to genes involved in Th17 pathogenicity (*Rbpj*, *Tbx21, Bhlhe40, Csf2, Ifng, Il17a*) ^6, 8, 33^ (GREAT analysis ^34^) as well as novel genes, including *Bach2*. Genes proximal to dLN-specific and CNS-specific DACRs were enriched for signatures previously identified from scRNA-seq of cells profiled from the dLN and CNS, respectively ^8^ (**Fig. 2D** and **Table S4**).

DACRs of dLN- and CNS-derived Th17 cells were enriched for motifs of putative regulators of non-pathogenic and pathogenic Th17 cells *in vivo* (**Fig. 2E**). dLN-specific peaks were enriched for sites for TFs associated with stem-like and self-renewing-like Th17 programs, including TCF1, LEF1, and OCT4 ^8, 16, 35^, whereas the CNS-specific peaks were enriched for sites for TFs known to play a role in Th17 differentiation and pathogenicity, including FOSL2, BATF, and JUNB ^36–38^.

To understand how the chromatin signatures from *in vitro* polarized cells related to chromatin accessibility changes that occur *in vivo* during EAE, we compared chromatin peaks accessible in both *in vitro*-derived and *in vivo*-derived Th17 cells (**Fig. 2F,G**). Because the CNS microenvironment differs greatly from *in vitro* polarization conditions and cells harvested *in vivo* represent a mixed ‘‘snapshot’’ of an asynchronous process with multiple heterogenous Th17 cell states ^8^, we focused only on 491 peaks with shared differential accessibility between npTh17 and dLN cells and 678 peaks shared between pTh17 and CNS cells (out of 41,168 peaks shared between *in vitro* and *in vivo* cells altogether, **Methods**). Peaks with shared differential accessibility in CNS-derived Th17 cells and pTh17 included loci involved in Th17 activation and pathogenicity (*Cd44*, *Stat4*, *Fosl2*, *Tbx21*, *Bhlhe40*) ^6, 8, 36^ as well as those not previously associated with Th17 cells (*Bach2*, *Ermn*). Differentially accessible peaks in both dLN-derived and npTh17 cells included loci characteristic of homeostatic and non-pathogenic Th17 cells (*Ccr6*, *Il6st*, *Maf*, *Il9*) ^6, 8^ and novel candidates (*Gpr15*, *Twist2*) (FDR <0.05). These peaks are consistent with the corresponding PC scores of ATAC-seq profiles *in vitro* and *in vivo*: those specifically accessible in npTh17 cells and dLN-derived Th17 cells with PC2 *in vitro*, and those shared in pTh17 cells and CNS-derived Th17 cells with PC1 *in vivo* (**Fig. 2F**, color bars).

Thus, chromatin accessibility profiles of Th17 cells from EAE-diseased mice revealed major differences between dLN-derived and CNS-derived Th17 cells in peaks near fate-defining genes, a subset of which is mirrored by *in vitro* generated npTh17 and pTh17 cells, respectively.

### The accessible chromatin features distinguishing pathogenic *vs*. non-pathogenic Th17 cells are shared with Th1 and T_reg_ cells, respectively

While Th17 cells are known to express genes typically attributed to other CD4^+^ T cell types (*e.g.*,Th1 genes *Ifng* and *Tbx21* expressed by pTh17 cells ^4, 12–15^), the relationship of Th17 cells to other T cell subsets and the extent of their plasticity have not been studied beyond selected markers. To better understand this, we next compared the ATAC-seq and RNA-seq profiles between *in vitro*-derived npTh17, pTh17, Th0, Th1, and T_reg_ cells (**Fig. 3A** and **S3A** and **Table S5, S6**).

**Fig. 3:**
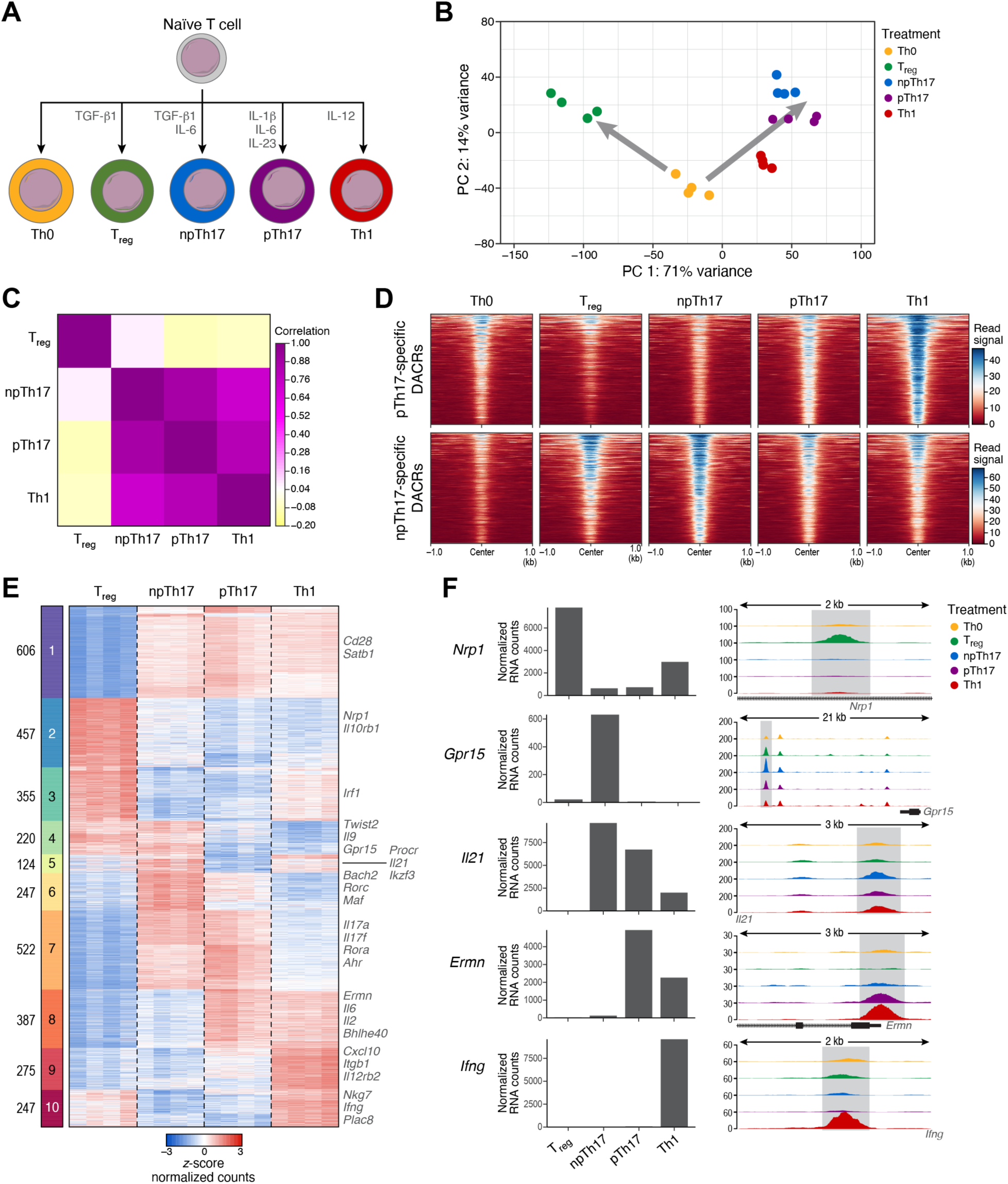
NpTh17 and pTh17 cells share different accessible chromatin features with other CD4^+^ T cells. (**A**) Experimental setup of *in vitro* T cell differentiation. Naïve CD4^+^ CD25^-^ CD44^-^ CD62L^+^ T cells were *in vitro* differentiated to Th0 cells (yellow), T_reg_ cells (green), npTh17 cells (blue), pTh17 cells (purple), or Th1 cells (red). (**B-D**) Unique and shared chromatin signatures define CD4^+^ T cell subsets. (**B**) First (*x* axis, PC1) and second (*y* axis, PC2) principal components of PCA of ATAC-seq profiles (dots) of *in vitro* differentiated Th0, npTh17, pTh17, Th1, and T_reg_ cells at 72h. Gray arrows denote two arms of CD4^+^ T cell polarization. (**C**) Pearson correlation coefficient (color bar) between pairs of normalized ATAC-seq peak count profiles of the CD4^+^ T cell subsets (rows, columns). (**D**) ATAC-seq normalized counts (color bars) in each CD4^+^ T cell subsets (panels) in the 1kb around the center of each peak of pTh17-specific (top rows) and npTh17-specific (bottom rows) DACRs, rank ordered by average signal in pTh17 and npTh17 cells, respectively. (E) Differential peaks in one or multiple different CD4^+^ T cell subsets. Z-score of normalized ATAC signal (color bar) in the replicate experiments in each cell type (columns) for the top 1,000 differentially accessible peaks between every pair of cell types (FDR <0.05, rows), ordered by *k*- means clustering (*k*=10, bar on left). Relevant genes associated with peaks are listed on the right. (F) CD4^+^ T cell-specific chromatin peaks near key marker genes. Gene expression (left, *y* axis, normalized counts) and enrichment of ATAC-seq signal over background (right, *y* axis, log fold) in the *Nrp1* (chr8:128,473,000-128,475,000), *Gpr15* (chr16:58,699,000-58,720,000), *Il21* (chr3:37,270,375-37,273,500), *Ermn* (chr2:58,051,000-58,053,517), and *Ifng* (chr10:118,406,000-118,408,011) loci in each CD4^+^ T cell subset (*x* axis on left, colored tracks on right). Gene bodies are displayed on the bottom. In cases of distal peaks the associated gene is indicated (right: downstream, left: upstream).

There were major differences between the CD4^+^ T cell subsets, with each subset grouping separately based on the first two PCs (**Fig. 3B** and **S3B,C**), forming two major “branches”, one from Th0 to Th1, pTh17 and npTh17 cells, and the other to T_reg_ cells. In both PCA and by pairwise correlation analysis, T_regs_ were the most distinct (**Fig. 3C**, Pearson’s *r* = 0.034, -0.133, -0.101 respectively for npTh17, pTh17, Th1 *vs*. Th0). While on PC1 (71% of variance) T_regs_ and npTh17cells were most distant (**Fig. 3B**), they were closest on PC2 (14% of variance), suggesting an underlying shared set of features. Indeed, this was also reflected in their RNA-seq profiles (**Fig. S3B**, PC2), possibly reflecting the shared requirement of TGF-β for their differentiation ^39^. Conversely, pTh17 cells were localized between the npTh17 and Th1 cells by both ATAC-seq (**Fig. 3B**) and RNA-seq (**Fig. S3B**), were most correlated to Th1 cells in their chromatin profiles (Pearson’s *r* = 0.810, 0.711, -0.101 Th1 *vs*. pTh17, npTh17, T_reg_, respectively, **Fig. 3C**), and DACRs that are more open in pTh17 *vs*. npTh17 cells displayed the highest signal in Th1 cells (**Fig. 3D**). NpTh17-specific DACRs were detected in T_reg_ cells (**Fig. 3D**), supporting that the dual functional cell states of pro-inflammatory and homeostatic Th17 cells reflect Th17 plasticity for other CD4^+^ T cell types.

To further compare chromatin landscapes across the Th cell subsets, we clustered the DACRs identified from pair-wise comparisons of npTh17, pTh17, Th1, and T_regs_ (*k*-means clustering; *k*=10, **Fig. 3E**), predicted genes associated with each DACR by proximity (using GREAT ^34^), and examined the expression of selected genes (from RNA-seq) (**Fig. 3F** and **S3C** and **S4**). Cluster 1, shared by all effector CD4^+^ T helper cells, included peaks close to *Cd28,* providing co-stimulatory signaling during T-cell activation ^40^, and *Satb1,* involved in the specification of CD4^+^ T cell subsets^41^. Cluster 2 and 3 consisted largely of T_reg_-specific peaks with some in proximity to genes important for T_reg_ function, such as *Nrp1*, *Il10rb1* and *Irf1* ^42–44^. Cluster 4 had DACRs shared between npTh17 and T_reg_ cells, including those near the genes *Twist2*, *Il9*, and *Gpr15*. IL-9 was previously implicated in Th17 differentiation and function and T_reg_-mediated immune suppression ^45–47^, and GPR15 is important for homing of T_reg_ cells to the large intestine ^32^. Cluster 5 and 6 harbored peaks specific to npTh17 cells, including some in proximity to genes previously associated with the npTh17 phenotype, such as *Procr*, *Il21*, *Ikzf3, Maf, Rorc* ^6, 8, 48^ and novel genes including *Bach2* (also associated by the *in vivo* analysis, **Fig. 2E,F**). Cluster 7 included peaks shared between both npTh17 and pTh17 cells and specific to the general Th17 phenotype, including those proximal to *Il17a*, *Il17f*, *Rora*, and *Ahr* genes ^36, 37^. Strikingly, cluster 8 highlighted peaks distinguishing pTh17 from npTh17 cells that are also accessible in Th1 cells, including near genes involved in proinflammatory and pathogenic Th17 responses such as *Bhlhe40* and *Il6* ^49–51^. Cluster 10 was specific to Th1 cells and included peaks close to Th1-marker genes, such as *Stat1*, *Il12rb2*, *Nkg7*, *Ifng*, and *Plac8* ^52–56^.

Overall, the npTh17 and pTh17 chromatin landscapes are distinct from those of other CD4^+^ T cell subsets through a set of shared DACRs and from each other through features that are shared with other Th cells – either T_regs_ (for npTh17) or Th1 cells (for pTh17), highlighting plasticity in Th17 cells that may drive differences in function.

### PTh17 cells retain an intermediate polarization state as npTh17 and Th1 cells diverge during a time course of polarization

To examine how the chromatin and expression changes at 48h arose over time we performed bulk ATAC-seq and bulk RNA-seq along *in vitro* polarization of Th1, pTh17 and npTh17 at 0hr, 1hr, 6hr, 12hr, 20hr, and 48hr (**Fig. 4A,B** and **S5A,B** and **Table S7**). In both ATAC-seq and RNA-seq, there were 3 main axes of variation by PCA: “time” (PC1), where samples from all three types progress similarly (**Fig. 4A,B** and **S5B**), “activation” (PC2), where all types increase from 1 to 6h (**Fig. S5B**), and “polarization condition” (PC3) (**Fig. 4A,B**), where samples from the different conditions grow increasingly distinct with time. In particular, samples from the pTh17 and npTh17 cells first both become distinct from Th1 cells (by 6h) and then npTh17 samples grow increasingly distinct from pTh17 cells, such that by 48h, pTh17 cells are an intermediate between Th1 and npTh17 cells. The similar patterns in ATAC-seq and RNA-seq **(Fig. 4A,B** and **S5B**) indicate that chromatin and expression changes are well-coordinated during T-cell polarization.

**Fig. 4:**
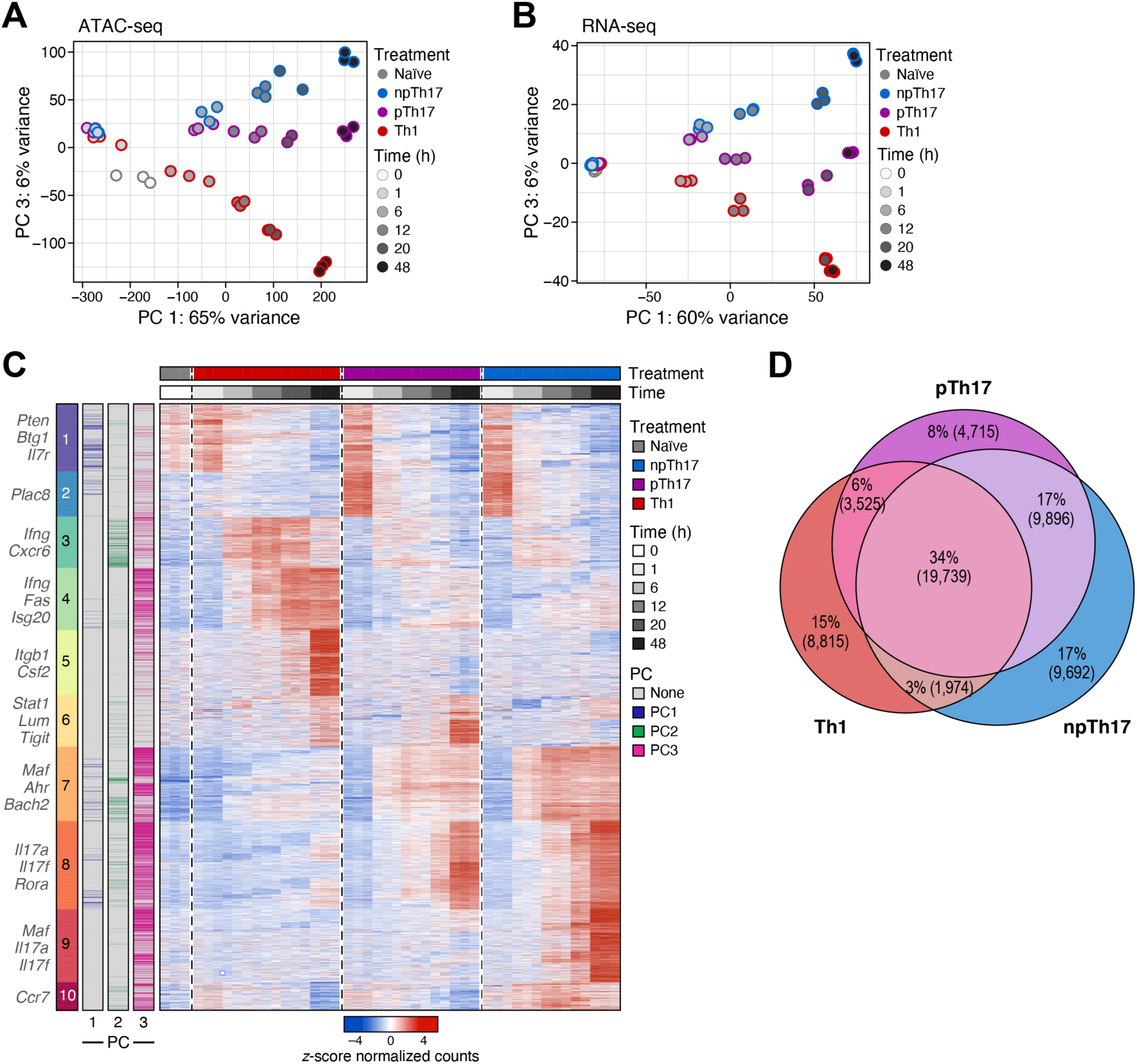
Divergence of npTh17 and pTh17 cells in early polarization leads to higher resemblance of pTh17 and Th1 cells. (**A,B**) Pathogenic Th17 cells occupy an intermediate chromatin and expression state between npTh17 and Th1 cells following polarization. First (*x* axis, PC1) and third (*y* axis, PC3) principal components of a PCA of bulk ATAC-seq (**A**) and bulk RNA-seq (**B**) of naïve and *in vitro* differentiated npTh17 (blue), pTh17 (purple) and Th1 (red) cells at 0h, 1h, 6h, 12h, 20h, and 48h (grey scale). (C) Changes in chromatin accessibility in npTh17, pTh17 and Th1 cells during polarization. Normalized counts (z-score, color bar) of the time dependent DACRs (FDR <0.05, rows) in Th1 (left, red), pTh17 (middle, purple), and npTh17 (right, blue) cells over time (greyscale bar, top), ordered by *k*-means clustering (*k*=10). Relevant genes are listed on the left. (D) Cell type specific and shared chromatin peaks that open during polarization. Chromatin regions that become more accessible in npTh17 (blue), pTh17 (purple) and Th1 (red) cells by 48h and their intersections.

Chromatin regions that were differentially accessible between the conditions over time partitioned into 10 clusters (**Fig. 4C**, *k*-means clustering, n = 10, 5,202 peaks), including cell type specific clusters (cluster 3, 4, 5, 9), and clusters that are shared between pTh17 and npTh17 cells (cluster 2, 7, 8), between pTh17 and Th1 cells (cluster 6), or across all conditions (cluster 1). Of the 5,202 top DACRs, 52% (2,708) were also in the top 20% peaks by loading from PC1, PC2, or PC3. Most (2,091 peaks, 77%) were shared with PC3, particularly in clusters 4, 5, 7, 8, 9, and 10, consistent with PC3 stratifying the polarization conditions (**Fig. 4C**, color bar). Of the 58,356 and 60,452 peaks that respectively opened or closed over the 48h, 34% (19,739) and 62% (37,204) were shared among the different conditions (**Fig. 4C,D** and **S5C**). Notably, in both the opening and closing peaks, the pTh17 condition shared more peaks with the npTh17 condition and the Th1 condition, than the Th1 condition shared with the npTh17 condition, further supporting that pTh17 cells are an intermediate state between npTh17 cells and Th1 cells.

### Individual pTh17 cells span an expression and chromatin spectrum between chromatin and expression features of Th1 and npTh17 cells

The shared programs and divergence patterns between pTh17 cells and Th1 or npTh17 cells could represent either a cell-intrinsic intermediate state or cell-to-cell heterogeneity within the pTh17 population (which would appear as an “intermediate” average bulk profile). To distinguish these possibilities, we next performed droplet-based scATAC-seq of *in vitro* differentiated npTh17, pTh17, and Th1 cells at 48h post-polarization.

ScATAC-seq profiles grouped by cell type (**Fig. 5A,B**, **S6A,B** and **S7A**), largely mirroring the Th1 to pTh17 to npTh17 order observed at the population level, suggesting a continuous, cell intrinsic phenotype (**Fig. 3B**). After excluding two small low-quality clusters (mixed, npTh17-2), the profiles partition in 5 clusters: one of npTh17 cells, two of pTh17 cells, and two of Th1 cells (**Fig. 5B**), each associated with different marker genes (by a gene score ^57^ of the ATAC-signal in a 200kb spanning window, **Fig. 5C**). The marker genes were consistent with those earlier identified with bulk ATAC-seq (**Fig. 1**) for the npTh17 (*Il17a*, *Il17f*, *Twist2*, *Il10*, *Havcr1*), pTh17 (*Themis*, *Il33*, *Fosl2*, *Il23r*) and Th1 (*Csf2*, *Ccl5*, *Tbx21*) clusters. Interestingly, the pTh17 cells spanned a spectrum (reflected by two “consecutive” clusters) from a higher chromatin accessibility at npTh17 marker genes (pTh17-2) to a higher shared chromatin program with Th1 cells (pTh17-1) (**Fig. 5B,C** and **S7B**). Accordingly, *in vivo* signatures of pathogenic Th17 cells were enriched in the pTh17-1 and Th1-1 clusters, while *in vivo* signatures of non-pathogenic Th17 cells were enriched in npTh17 and pTh17-2 clusters (**Fig. S7C,D**).

**Fig. 5:**
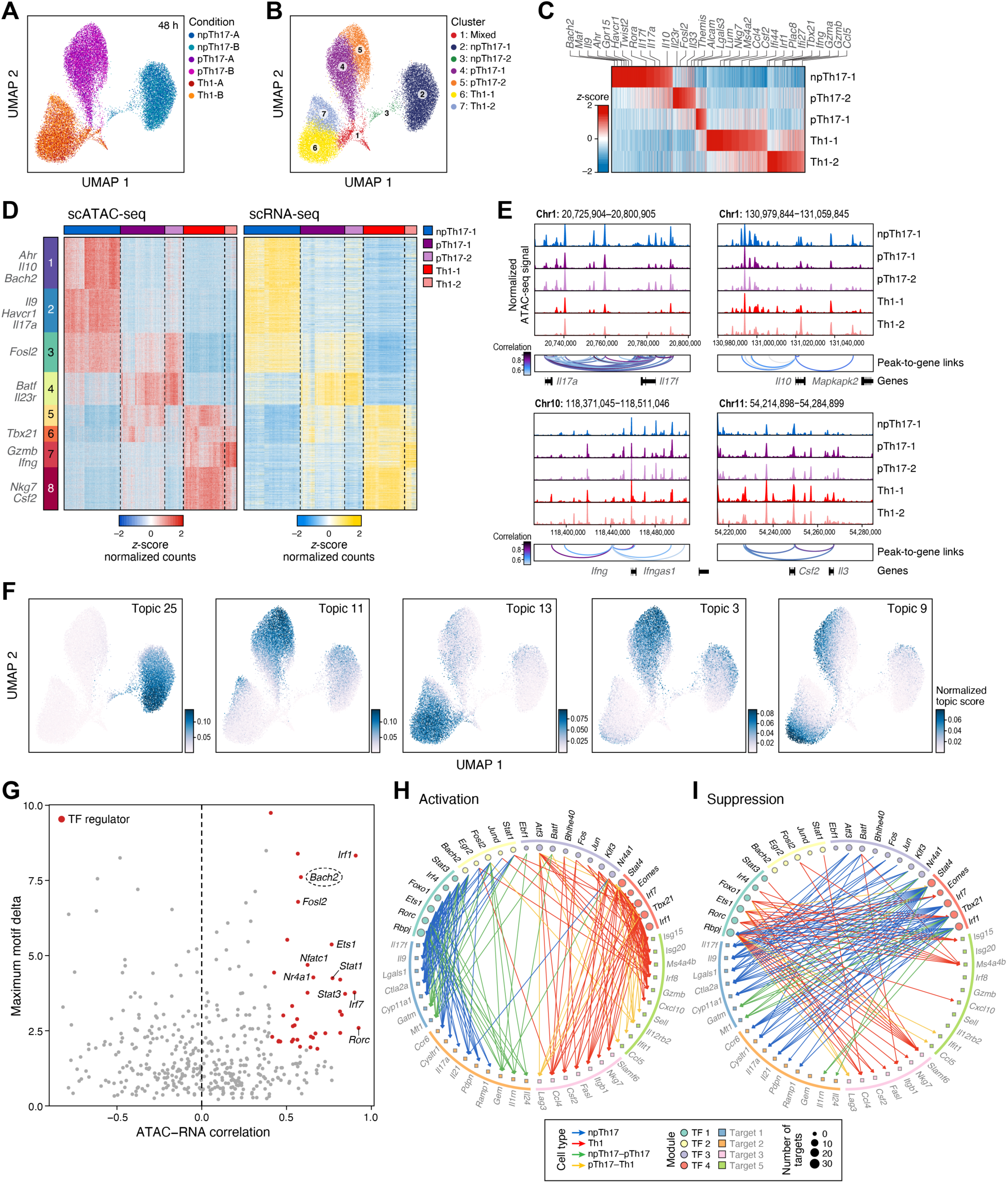
Single-cell chromatin accessibility and RNA profiling reveal a spectrum of chromatin profiles of pTh17 cells and enable TF regulator discovery. (**A-C**) Heterogeneity in scATAC-seq profiles of npTh17, pTh17, and Th1 cells at 48h. (**A,B**) Uniform Manifold Approximation and Projection (UMAP) embedding of scATAC-seq profiles (dots) colored by cell subset (**A**) or by cluster membership (**B**). (**C**) ATAC-seq signal (gene scores; z-score, color bar) at loci of marker genes (columns, 100kb on either side of the gene) for each cluster (rows). (**D,E**) Peak accessibility and gene expression are correlated across Th17 cells. (**D**) scATAC-seq (left) and scRNA-seq (right) signals (z-score of normalized counts) of peaks (left rows) and genes (right rows) related by peak-to-gene correlation. Peaks (left) and their related genes (right) are both ordered by *k*-means clustering of ATAC-seq peak profiles (*k*=8). Relevant genes are listed on the left. (**E**) Normalized ATAC-seq signal (*y* axis) at selected Th17 gene loci showing peak-to-gene links (bottom arcs, colored by peak-to-gene correlation, color bar). (F) Distinct and shared gene programs distinct across Th17 and Th1 cells identified by topic modeling. UMAP embedding of scATAC-seq profiles (dots) colored by topic scores per cell (normalized topic score, colorbar) for selected topics. (G) Inferred positive regulators of the Th17 chromatin landscape. Maximum motif delta (*y* axis), defined as the highest TF motif deviation score driving variation between clusters, and the correlation of the corresponding TF’s gene chromatin accessibility score with the motif deviation (*x* axis) across single cells. Red: significant positive regulators with a correlation >0.4 and maximum motif delta >4 (FDR <0.05). BACH2 is circled. (**H,I**) Inferred positive (**H**) and negative (**I**) regulatory interactions in a Th17 and Th1 regulatory subnetwork. A directed edge is shown from a TF (circles, top) to its predicted positively (**H**) and negatively (**I**) correlated target gene (squares, bottom). Edge color: cell type(s) in which the association was detected (color legend). Only a selected subset of genes and their interactions are shown.

To link accessible chromatin regions to the potential target genes they regulate, we also profiled *in vitro* polarized npTh17, pTh17, and Th1 cells by scRNA-seq, observing a similar ordering of Th1-pTh17-npTh17 profiles (**Fig. S7E,F**). We integrated scATAC-seq and scRNA-seq profiles, assigned RNA profiles to each cell from our scATAC-seq, and used these to correlate chromatin peak accessibility with gene expression ^57^ (**Fig. 5D,E** and **Table S8**). The 23,608 identified peak-to-gene links (correlation >0.45, FDR <0.05) (**Fig. 5D**) partitioned to eight clusters (*k*-means clustering). These included both cell type-specific peak clusters for npTh17 cells (cluster 1, 2; 8,260 peak-to-gene links, *e.g.*, *Ahr*, *Il10*, *Bach2*, *Il9*, *Havcr1*, *Il17a*), pTh17 cells (cluster 4, 2,827 links, *e.g.*, *Batf*, *Il23r*), and Th1 cells (cluster 7, 8, 5,888 links, *e.g.*, *Gzmb*, *Ifng*, *Nkg7*, *Csf2*, **Fig. 5D**), as well as a shared npTh17 and pTh17 cluster (cluster 3, 3,442 links, *Fosl2*) and shared pTh17 and Th1 clusters (clusters 5, 6, 3, 191 links, *Tbx21*). The peak-to-gene links included key Th17 effector gene loci, such as *Il17a*, *Il17f*, *Il10*, *Csf2*, and *Ifng*, with peaks that highly correlated with gene expression and may act as potential regulatory elements (**Fig. 5E**).

Consistent with our scATAC-seq analysis, cells in the pTh17-2 scRNA-seq cluster had higher signal for peaks shared with npTh17 cells, while pTh17-1 cluster cells had more accessible chromatin in clusters 5 and 6 that contain pro-inflammatory and Th1-specific genes. Notably, in the peak-to-gene linkage, a given gene can be correlated with multiple peaks (**Fig. 5E**). Multiple peaks associated with pTh17 gene markers (*Il23r, Il22, Themis)* were all highly specific to the same cell-type and thus members of the same single cluster. Conversely, genes associated with npTh17 phenotypes (*Il17a, Il10, Ahr, Bach2)* and pro-inflammatory Th1-like states (*Tbx21, Ifng, Gzmb, Tbx21*) were matched to peaks found in different cell-type specific clusters, revealing how these genes may be regulated in a cellular context-dependent manner (**Fig. S7G-I**).

Topic modeling on scATAC-seq data ^58^ captured 35 cellular programs and states (“topics”) some distinguishing one or two of Th1, pTh17 and npTh17 cells, others varying within a single cell subset, and yet others varying within two of the subsets in similar patterns (**Fig. 5F** and **S8A**). In the cell type specific topics, Topic 25 featured accessible chromatin regions defining npTh17 cells, including peaks linked to npTh17 marker genes *Il17a, Batf, Il10, Twist2,* and *Rbpj*; Topic 11 distinguished pTh17 cells with accessible chromatin regions proximal to pathogenicity-associated genes such as *Il23r*, *Il21*, and *Bhlhe40*; and Topic 13 represented a Th1 program with accessible chromatin regions associated with pro-inflammatory gene programs (*Csf2, Eomes, Fas, Cd44*) (**Fig. 5F** and **S8B**). Notably, even such specific topics scored in minor portions of the other cell types, showing the malleability of these programs. This malleability was even more striking in other topics that captured variation *within* multiple cell types, such as a shared npTh17 and pTh17 program (Topic 3), with peaks proximal to *Il17a, Il17f, Satb1*, and *Bach2*, and a shared pTh17-1 and Th1-1 program (Topic 9) in pTh17-1 and Th1-1 clusters, defined by peaks near pro-inflammatory genes *Il12a, Il12rb2, Atf3, Il33*, and *Ccl5* (**Fig. 5F** and **S8B**).

Overall, this analysis showed that pTh17 cells span a spectrum, from cells where npTh17-like features are more prominent to cells with a stronger pro-inflammatory Th1-like chromatin state.

### Combined scATAC-seq and scRNA-seq profiles recover a self-reinforcing, mutually antagonistic regulatory network across npTh17, pTh17, and Th1 cells

Next, we used the integrated scATAC-seq and scRNA-seq profiles to predict putative TF regulators that bind at T-cell type-specific accessible chromatin sites, highlighting both known regulators of Th17 cells and novel candidates. To this end, we first identified TFs whose binding site motifs are enriched in Th1, npTh17, and pTh17-specific accessible chromatin regions (**Fig. S9A**). Next, we further identified TFs whose own RNA expression positively correlated with changes in the accessibility of regions with their corresponding motif across loci, by comparing the integrated RNA expression of a given TF to the TF motif deviation ^59^ (**Fig. 5G** and **Table S9**). This analysis highly ranked known regulators of Th17 cells including *Ets1* ^60^, *Junb*, and *Fosl2* ^36–38^, as well as a novel candidate regulator, the transcription factor BTB Domain and CNC Homolog 2 (BACH2).

We further leveraged our matched scATAC-seq and scRNA-seq data to construct a TF:target regulatory network across the Th17 and Th1 programs. We identified differentially expressed genes from npTh17, pTh17, and Th1 scRNA-seq (FDR <0.01, fold change >2), and then searched for TF motifs in the peaks that were associated with these genes by peak-to-gene linkage analysis. The resulting network related a TF to a target gene if the TF motif was found in the peak linked to the gene and the TF and target gene expression were correlated across all cells in scRNA-seq (Pearson |*r*| >0.1, FDR <0.001). Focusing on the highly interactive TF regulators (with at least 10 targets) and target genes (of at least 5 TFs), the resulting network had 78 TFs, 223 target genes (including 15 that were also TF regulators), and 3,749 edges in the full network (**Fig. S9B**). Within it, a focused subnetwork (**Fig. 5H,I**) spanned 24 TFs and 33 target genes with 201 activation (**Fig. 5H**) and 114 suppression (**Fig. 5I**) edges. Several of the network TFs, including *Bach2*, *Fosl2*, *Irf1*, *Ets1, Nr4a1* and *Rorc*, were also predicted as top regulators of chromatin accessibility (**Fig. 5G**).

The network architecture suggests a largely self-reinforcing, mutually antagonistic organization, such that “activators” of gene programs for one state act as repressors for the other, and vice versa, including regulation between the TFs (**Fig. 5H,I** and **S9B**). Specifically, hierarchical clustering of the TF:target expression correlation matrix (**Fig. 5H,I** and **Fig. S9B**) recovered 4 TF modules and 5 target gene modules (**Fig. S9B**). TF modules 1 and 2 positively correlated with target modules 1 and 2 in npTh17 and pTh17 cells (**Fig. 5H,I** and **S9B**, blue and green edges). TF module 1 (17 TFs) and 2 (24 TFs) included established master TFs of the Th17 cell fate, such as *Rorc*, *Irf4*, *Stat3*, *Batf3,* and *Fosl2* ^36, 37^ and the novel candidate regulator *Bach2*. Target module 1 (36 genes) and 2 (64 genes) included Th17-marker genes (*Il17f* and *Il9* in module 1*; Ccr6*, *Il17a*, *Il21*, and *Pdpn* in module 2). TF modules 3 and 4 positively correlated with the pTh17-Th1 target modules 3 (50 genes) and 5 (47 genes), which included known pathogenicity genes (*e.g., Nkg7* ^61^, *Ccl5* ^62^, *Fasl* ^63^, *Csf2* ^16^). The smallest target module 4 (11 genes) included genes specifically expressed in pTh17 cells and did not show a strong correlation with any of the TF modules. At the same time, each of the TF module pairs negatively correlated with the target modules: TF modules 1 and 2 with target modules 3 and 5 (in pTh17 and Th1 cells) and TF modules 3 and 4 with target modules 1 and 2 (in npTh17 and pTh17 cells). In these contrasting roles of TFs in the network, positively correlating with target genes from npTh17 or Th1 cells but negatively correlating with genes from the other subset, is consistent with previous network architectures we have observed in Th17 cells at the expression level ^37^. Notably, this pattern was also observed when considering only TFs as targets, consistent with a self-reinforcing, mutually exclusive regulatory architecture.

### Network analysis predicts regulators of Th17 cell pathogenicity

In line with our previous findings, pTh17 TFs and genes were primarily in modules that are shared with either npTh17 or Th1, highlighting the regulatory architecture underlying our finding that pTh17 cells form a state within a spectrum that is intermediate between npTh17s and Th1s, and shared distinct features with each. For example, the pTh17-Th1 target modules 3 and 5 (**Fig. 5H** and **S9B**) are regulated by *Tbx21*, *Eomes*, *Nr4a1*, and *Atf3*. *Eomes* and *Nr4a1* modulate Th17 and Th1 fates in autoimmunity and inflammation ^64, 65^. ATF3 regulates *Ifng* in Th1 cells ^66^ and *Il10* in Tr1 cells ^67^, but its roles in driving Th17 pathogenicity have not yet been described. Similarly, *Rorc*, *Ets1*, *Fosl2*, *Rbpj,* and *Bach2* are among the TFs that regulate the npTh17-pTh17 target modules 1 and 2 (**Fig. S9B Fig. 5H**). *Rbpj* is a known regulator of Th17 gene programs ^33^ and is identified here as a central regulator (168 target genes, **Fig. S9B**). *Bach2* (46 target genes, **Fig. S9B**) has not been implicated in Th17 gene regulation but is predicted to be a top driver of the Th17 gene programs in this model.

Many of the regulators identified by network analysis at 48h are also identified by TF motif enrichment analysis on the peaks of each cluster of chromatin regions that change along the polarization time course (**Fig. 4** and **Fig. S9C**). For example, peaks in the Th1-specific cluster (cluster 4) are enriched with motifs of TFs involved in Th1 differentiation, NFAT, NF-κB, and NUR77 ^65, 68–70^. Cluster 7 included chromatin regions common in Th17 cells that opened early (6h) and were predicted to be regulated by known Th17 regulators, including BATF, JUNB, and FOSL2^36, 38^ and the novel regulators ATF3 and BACH2. Cluster 8 included Th17-specific peaks that appeared later during differentiation (20-48h) and were enriched with motifs of the known Th17 regulators BATF, JUNB and RORγt ^36–38, 71^, indicating that the same TFs regulate two different waves of Th17-specific peaks during Th17 differentiation.

In summary, ATAC-seq and RNA-seq at the single cell level and across multiple time points of Th17 and Th1 differentiation allowed us to predict many known regulators of the Th17 chromatin landscape, organized in a self-reinforcing, mutually exclusive architecture across the cell programs, as well as novel ones, such as the TF BACH2, associated with npTh17 programs.

### BACH2 is a negative regulator of Th17 pathogenicity by restraining Th1 programs in Th17 cells

We next aimed to functionally validate BACH2, which emerged as a novel regulator of the Th17 chromatin landscape predicted from our analysis across both *in vitro* and *in vivo* data. Multiple lines of evidence support *Bach2* as putative driver of Th17 cell states: (**1**) BACH2 was a highly ranked candidate in the TF motif analysis of peaks more accessible in the CNS than in the dLN (**Fig. 2E**); (**2**) ATAC-seq peaks close to the *Bach2* locus were more open in Th17 cells from the CNS than from the dLN (**Fig. 2F**); (**3**) npTh17-specific peaks were found in proximity to the *Bach2* locus (**Fig. 3E**); (**4**) BACH2 is a top positive regulator of npTh17 targets based on integrated scATAC-seq and scRNA-seq (**Fig. 5G,H**); and (**6**) BACH2 was predicted to regulate Th17- specific chromatin regions over time, especially in npTh17 conditions (**Fig. S9C**). Notably, although the *Bach2* motif was highly enriched in DACR peaks specific to CNS-derived Th17 cells, *Bach2* gene expression in the bulk RNA-seq was only marginally higher in CNS Th17 cells compared to dLN-derived Th17 cells (log2FC = 0.52, FDR = 0.25), highlighting the benefit of motif information derived from chromatin accessibility when searching for transcriptional regulators.

To test our hypothesis, we used the CRISPR/Cas9 system to create loss-of-function mutations (LOF) in the protein-coding region of *Bach2*. We obtained naïve T cells from Cas9 knock-in mice ^72^ and *in vitro* differentiated them to npTh17 and pTh17 cells while simultaneously transducing them with retroviral vectors encoding a *Bach2*-targeting single guide RNA (sgRNA) (*Bach2* KO) or empty sgRNA as control. We confirmed highly efficient indel formation at the target *Bach2* locus in Th17 cells (**Fig. S10A**).

Knockout (KO) of *Bach2* during Th17 differentiation led to increased expression of the pro-inflammatory cytokines IL-17A (p <0.05, unpaired t-test), IFN𝛾 (p <0.001), and GM-CSF (p <0.01) in npTh17 cells (**Fig. 6A**), suggesting that BACH2 restrains Th17 pathogenicity in npTh17 cells.

**Fig. 6:**
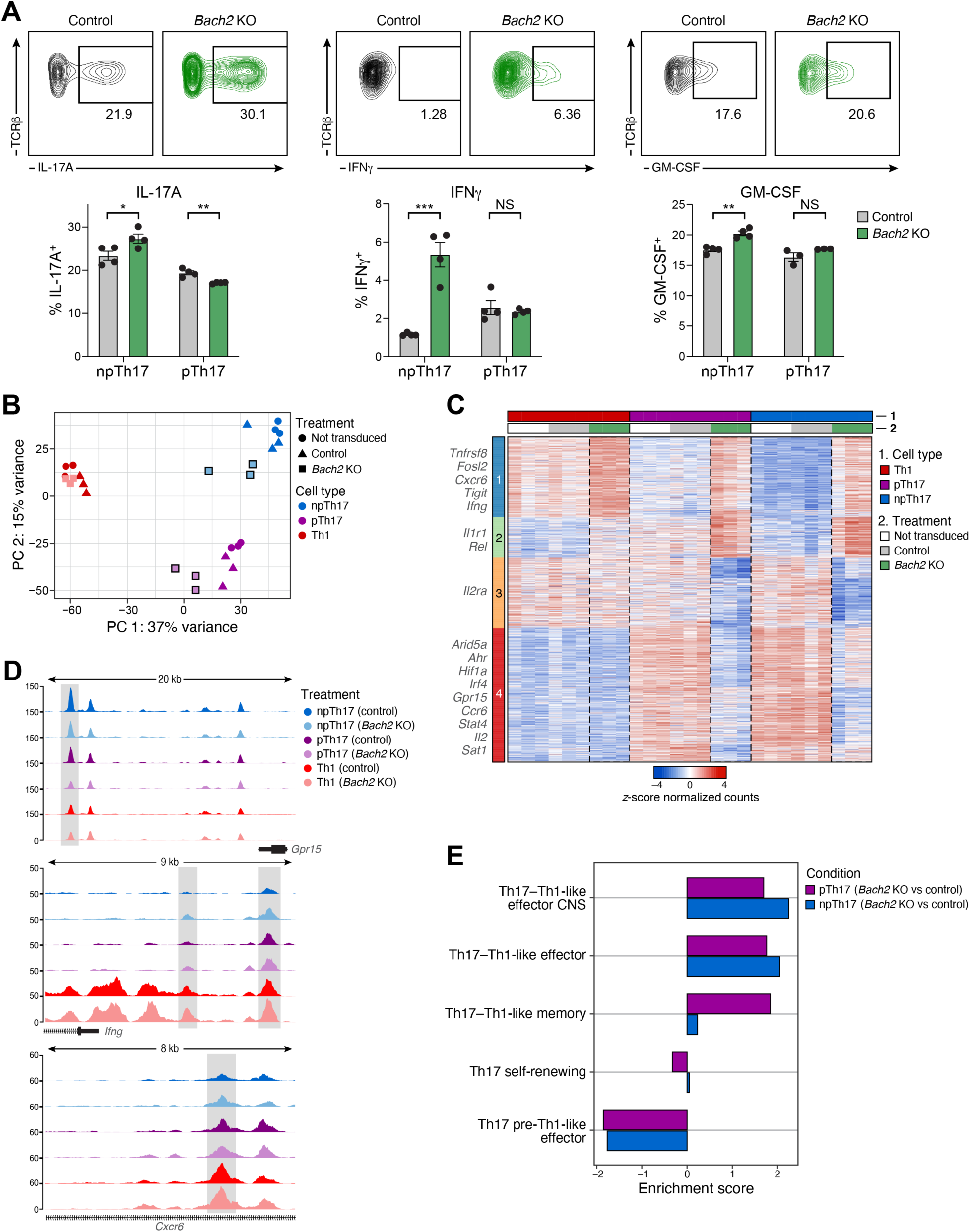
BACH2 restrains Th1 cell fate programs in npTh17 and pTh17 cells. (**A**) *Bach2* KO increases pro-inflammatory cytokine expression in npTh17 cells. Percent (*y* axis bottom, mean with ± SEM) of npTh17 and pTh17 cells (*x* axis, bottom) expressing IL-17A (left), IFNγ (middle), and GM-CSF (right) by flow cytometry (illustrative top panels) in Th17 cells with retroviral transduction of a *Bach2*-targeting gRNA (*Bach2* KO, green) or a non-targeting gRNA (controls, grey). *, P <0.05; **, P <0.01; ***, P <0.001, NS, not significant, unpaired two-tailed t-test. (**B-D**) *Bach2* KO shifts npTh17 and pTh17 cells toward a pro-inflammatory Th1-like state. (**B**) First (*x* axis, PC1) and second (*y* axis, PC2) principal components of a PCA of bulk ATAC-seq of *Bach2* KO (squares), control (triangles), and untransduced (circles) cells in npTh17-, pTh17-, and Th1- differentiation. (**C**) ATAC-seq signal (z-score) in 1,476 DACRs (rows) that are differentially accessible (FDR <0.05, |log2(fold change)| >0.5) between *Bach2* KO (green), control (grey) and not transduced (white) conditions, and clustered by *k*-means clustering (*k*=4). Peaks shown are the top 500 DACRs by positive log_2_(fold change) and bottom 500 DACRs by negative log_2_(fold change) comparing *Bach2* KO *vs.* control in each cell type. Relevant genes associated with the ATAC-peaks are listed on the left. (**D**) Enrichment of ATAC-seq signal over background (log fold, *y* axis) in the *Bach2* KO (dark color) and control (light color) conditions in npTh17 (blue), pTh17 (purple) and Th1 (red) cells at the *Gpr15* (chr16:58,700,000-58,719,516), *Ifng* (chr10:118,443,974-118,452,930), and *Cxcr6* (chr9:123,808,813-123,817,000) loci. Gene bodies are displayed on the bottom. (**E**) Differentially accessible loci in *Bach2* KO are enriched for pathogenic Th17 signatures. Enrichment (*x* axis) of Th17 pathway signatures ^8^ (*y* axis) in genes associated with the differential ATAC-peaks in *Bach2* KO *vs.* control in npTh17 cells (purple, 1,118 peaks) and in pTh17 cells (blue, 627 peaks) (**Table S11**).

BACH2 substantially impacted the chromatin landscape of Th17 cells but had little effect on Th1 cells, based on bulk ATAC-seq of *Bach2* KO and control npTh17, pTh17, and Th1 cells, shifting both pTh17 and npTh17 cells more towards a Th1 chromatin profile, suggesting that BACH2 restricts the Th1 program in Th17 cells (**Fig. 6B-D** and **Table S10**). Specifically, while *Bach2* KO in Th1 cells resulted in only modest chromatin accessibility changes (18 DACRs), the *Bach2* KO npTh17 cells (1,118 DACRs) and pTh17 cells (627 DACRs) demonstrated substantial changes (FDR <0.05, |log2(fold change)| >0.5). Moreover, *Bach2* KO profiles from both npTh17 and pTh17 cells are shifted toward Th1 cells on PC1 (**Fig. 6B**), with opening of Th1-specific peaks in proximity to the genes *Ifng, Fosl2*, and *Tigit*, and closing of Th17-specific peaks in proximity to the genes *Ahr*, *Gpr15*, and *Stat4* (**Fig. 6C,D**). In addition, there was Th17-specific closing of peaks in the *Bach2* KO in the *Il2ra* locus and opening of a set of peaks in the *Il1r1* locus in all *Bach2* KO cells (**Fig. 6C**). Accordingly, genes proximal to *Bach2* KO-specific peaks in pTh17 and npTh17 cells were enriched for *in vivo* signatures of Th17 pathogenicity and depleted for signatures of non-pathogenic Th17 phenotype ^8^ (**Fig. 6E** and **Table S11**). Moreover, chromatin accessibility signatures of T_reg_ and Th17 cells (**Fig. 3E**) are down-regulated in both npTh17 and pTh17 cells from *Bach2* KO (**Fig. S10B**).

Consistent with the KO phenotype, *Bach2* overexpression (OE) in pTh17 cells polarized *in vitro* led to closing of Th1-like pathogenic features, suggesting it acts as a repressor (**Fig. 7A,B**). Specifically, we polarized naïve CD4^+^ T cells into pTh17 cells by simultaneous transduction of retroviral vectors with a constitutive *Bach2* cDNA expression cassette (*Bach2* OE) or an empty control and achieved a significant increase (∼10-fold, p <10^-4^, unpaired t-test) in *Bach2* expression compared to control (**Fig. S11A**). We profiled *Bach2* OE and control populations by bulk ATAC-seq. *Bach2* OE led to major changes in the chromatin landscape of pTh17 cells (**Fig. 7A,B** and **Table S12**), opening 524 peaks, including close to genes associated with the stem-like, non-pathogenic Th17 phenotype (*Ccr7*, *Maf*, *Tcf7*), and closing many more peaks (3,438 peaks), including in proximity to genes involved in Th17 pathogenicity, such as *Bhlhe40*, *Csf2*, *Cxcr6*, *Ermn*, *Ifng*, and *Rbpj* ^6, 8, 16, 33^.

**Fig. 7:**
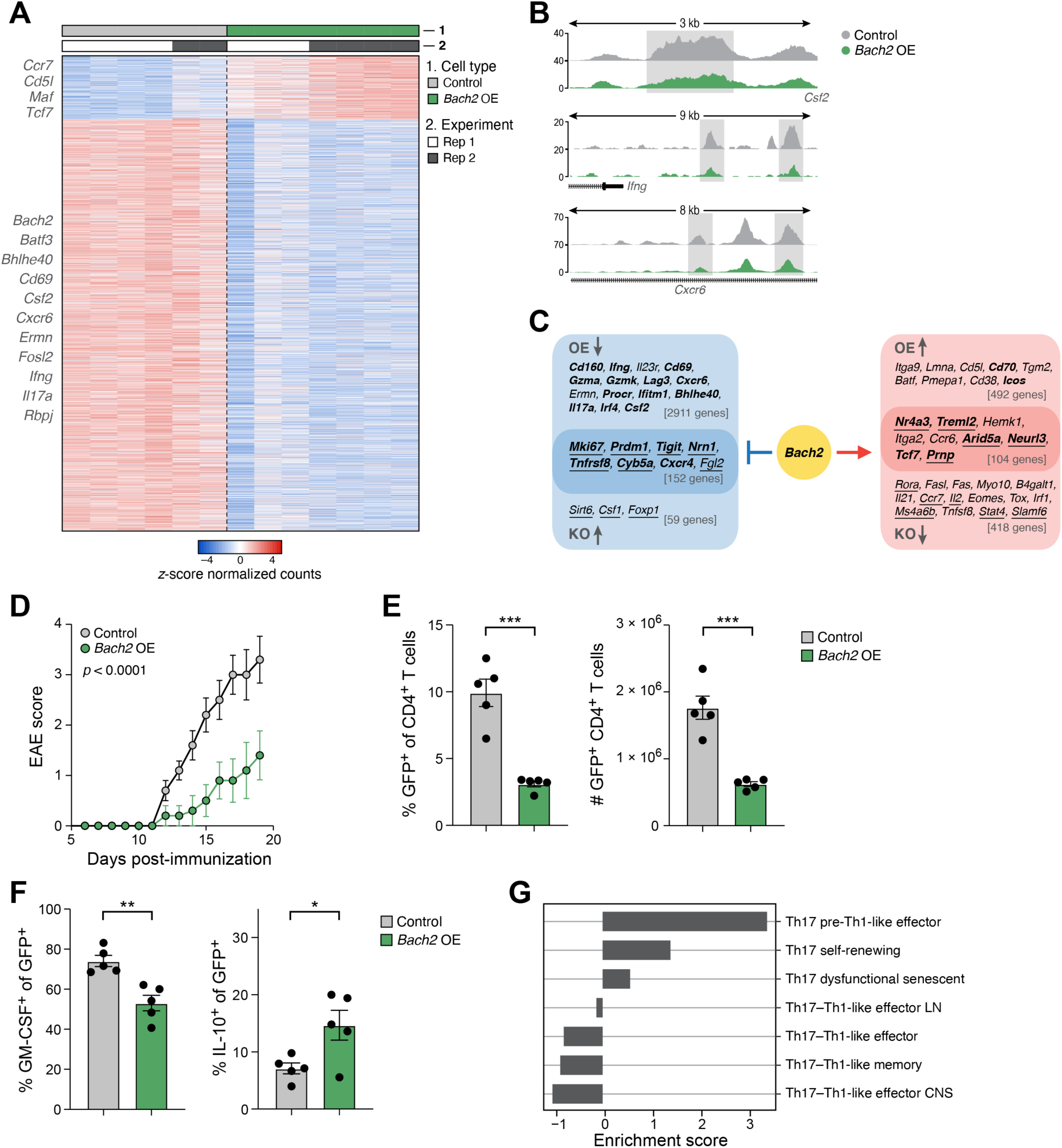
*Bach2* overexpression diminishes Th17 pathogenicity. (**A,B**) *Bach2* OE in pTh17 cells reduces chromatin accessibility at pro-inflammatory genes. (**A**) Relative ATAC signal (z-score of normalized counts, color bar) in ATAC-seq peaks (rows) that are differentially accessible (FDR <0.05, |log2(fold change)| >0.5) between *Bach2* OE (green columns) and control (grey columns) mice. Relevant genes associated with the peaks are highlighted on the left. (**B**) Enrichment of ATAC-seq signal over background (*y* axis, log fold) in the *Bach2* OE (green) and the control (grey) mice in the *Csf2* (top, chr11:54,213,319-54,216,630), *Ifng* (middle, chr10:118,443,974-118,452,930), *Cxcr6* (bottom, chr9:123,808,813-123,817,000) loci. Gene bodies are displayed on the bottom. In cases of distal peaks the associated gene is indicated (right: downstream). (**C**) Key genes regulated by BACH2. Schematic of genes inferred as repressed (left) or activated (right) by Bach2 based on their respective association with differential accessibility in *Bach2* OE or *Bach2* KO *vs.* the respective control (FDR <0.05 and |log2(fold change)| >0.5; genes showing contradictory directions in OE and KO experiments were excluded). Bolded (underlined) gene names: genes with a *Bach2* motif in the peak associated with *Bach2* OE (KO). (**D-F**) *Bach2* OE 2D2 cells transfer diminished EAE disease. (**C**) Clinical EAE score (*y* axis, mean with ± SEM) over time (*x* axis) of mice adoptively transferred with *Bach2* OE (green) or control (grey) 2D2 cells. (**E,F**) Frequency (**D**, left) and number (**D**, right) of transduced (GFP^+^) cells and expression of GM-CSF (**E**, left) and IL-10 **(E**, right) in transduced cells (GFP^+^) in the CNS of EAE recipients of *Bach2* OE (green) or control (grey) pTh17 2D2 cells by flow cytometry. Data are mean ± SEM. *, P <0.05; **, P <0.01; ***, P <0.001, unpaired two-tailed t-test. n=5 mice. (**G**) Enrichment score (*x* axis) of Th17 pathway signatures ^8^ (*y* axis) in differentially expressed genes from bulk RNA-seq of transduced (GFP^+^) *Bach2* OE *vs.* control 2D2 cells isolated from the CNS (**Table S13**).

The changes in chromatin accessibility in *Bach2* CRISPR KO and *Bach2* OE in polarized Th17 cells are often diametrically opposed. DACRs in 104 genes became more closed in Bach2 KO and more open in Bach2 OE compared to their controls (418 genes in *Bach2* KO alone and 492 genes in *Bach2* OE alone) (**Fig. 7C**). DACRs in 121 genes became more open in *Bach2* KO and more closed in *Bach2* OE (59 genes in *Bach2* KO alone and 2,911 genes in *Bach2* OE alone, **Fig. 7C**). Many of these target genes are potentially directly bound by BACH2, as indicated by the presence of a BACH2 binding motif found in the associated DACR, including stem-like T cell marker *Tcf7* and *Slamf6* as well as pro-inflammatory genes *Ifng* and *Csf2*. In *Bach2* KO, 147 of 211 (69.7%) gene peaks that became more open and 95 of 522 (18.2%) gene peaks that became more closed contained a BACH2 binding motif (p = 1 x 10^-20^ for *Bach2* motif enrichment in opened *vs.* closed peaks). In *Bach2* OE, a BACH2 binding motif was detected in 1,370 of 3,063 (44.7%) gene peaks that became more closed and 128 of 596 (21.5%) gene peaks that became more open (p = 1 x 10^-104^ for *Bach2* motif enrichment in closed *vs.* open peaks). This is consistent with the reported action of BACH2 as a transcriptional repressor ^73, 74^.

Thus, KO of *Bach2* in Th17 cells leads to an upregulation of a pathogenic Th1-like chromatin program in both npTh17 and pTh17 cells, and its OE in pTh17 cells leads to downregulation of this program (and induction of a stem-like non-pathogenic Th17 phenotype). BACH2 may thereby act as a repressor, restraining Th1-like features and Th17 pathogenicity in Th17 cells during differentiation.

### Increased expression of BACH2 ameliorates Th17 autoimmunity *in vivo*

Given the role of BACH2 in restraining chromatin features of Th17 pathogenicity *in vitro* (**Fig. 6** and **7A,B**), we hypothesized that BACH2 overexpression in pTh17 cells could ameliorate EAE disease *in vivo*. To test this hypothesis, we used the 2D2 EAE transfer model, in which the adoptive transfer of CD4^+^ T cells from 2D2 T-cell receptor (TCR) transgenic mice that express a TCR specific for the myelin oligodendrocyte glycoprotein (MOG) ^75^ induces severe paralytic disease in the recipient mice. We *in vitro* polarized naïve CD4^+^ T cells to pathogenic *Bach2* OE or control 2D2 Th17 cells (as above) and adoptively transferred them into congenically-marked recipient mice.

While the transfer of T cells treated with the control virus induced severe EAE disease, transfer of T cells overexpressing *Bach2* resulted in significantly milder disease (**Fig. 7D**, p <10^-4^) and had significantly lower frequencies (p <10^-3^, unpaired t-test) and numbers (p <10^-3^, unpaired t-test) of transduced (GFP^+^) T cells in the CNS (**Fig. 7E**), and those cells expressed the pathogenic cytokine GM-CSF at significantly lower frequencies (p <0.01, unpaired t-test) and the regulatory cytokine IL-10 at higher frequencies (p <0.05, unpaired t-test) (**Fig. 7F**). Bulk RNA-seq of the transduced (GFP^+^) cells from the CNS of EAE-diseased mice showed that the expression profile of *Bach2* OE cells was enriched for signatures of *in vivo* non-pathogenic and stem-like Th17 cells ^8, 16^, while the control cells exhibited a highly pathogenic profile with signatures of Th17, pre-Th1 effector like cells (**Fig. 7G** and **S11B** and **Table S13**). Moreover, an *in vivo* stem-like Th17 signature ^16^ was enriched in *Bach2* OE cells and down-regulated in *Bach2* KO cells (**Fig. S11C**). This is consistent with a recent study showing that BACH2 is a regulator of the stem-like program in CD8^+^ T cells ^76^, and further suggests BACH2 as a driver of a stem-like expression program in T cells more broadly.

Thus, the ameliorated *in vivo* pathogenicity of *Bach2* overexpressing pTh17 cells validates *Bach2* as a novel regulator of the Th17 pathogenic chromatin landscape and suggests *Bach2* as a potential target for the treatment of autoimmune conditions.

### BACH2 variants are associated with autoimmune disease risk in humans

Finally, we examined a potential association of BACH2 with autoimmune disease in humans. In the genome-wide association study (GWAS) of MS ^77^, *BACH2* is prioritized as a putative causal gene associated with a protective GWAS locus tagged by rs72928038 (odds ratio for G allele: 0.8663; p = 8.38*10^-29^) (**Fig. S12A,B**). Fine mapping of the locus reports three variants that explain the GWAS signal: rs72928038 (posterior probability: 84.2%), rs10944479 (posterior probability: 10.4%), and rs6908626 (posterior probability: 1.1%) ^78^.

Chromatin accessibility, chromosome conformation and expression data further suggest that the most strongly associated variant, rs72928038, is located in a regulatory element whose function could be linked to *BACH2* transcriptional regulation. First, rs72928038 resides in an open chromatin peak intronic on *BACH2* that is open in all primary immune cells ^78^, suggesting potential for regulation of nearby genes. Moreover, promoter capture interaction data (PChiC) ^79^ shows an interaction between the rs72928038 locus and the *BACH2* promoter in all 17 examined cell types and states, but only reaching study-wide significance (by FDR) in naïve CD4^+^ T cells (interaction score = 9.28) (**Fig. S12C**). Next, publicly available eQTL data (DICE consortium ^80^) showed that rs72928038 and rs10944479, the top two of the three putative causal variants, were associated only with changes of expression of *BACH2*. Carriers of the rs72928038 G allele had higher *BACH2* expression in naïve CD4^+^ T cells (adjusted p = 0.0015) and in T_reg_ cells (adjusted p = 0.0018) (**Fig. S12D,E**). These results suggest that the protective allele for MS susceptibility can result in increased *BACH2* expression in CD4^+^ T cells, in agreement with our results in mouse Th17 cells.

## Discussion

In this study, we leveraged the accessible chromatin landscape of CD4^+^ T cells along with RNA profiles in bulk populations and single cells to understand the distinctive cell fate features and relationships between Th17 cells, pTh17 cells and other CD4^+^ T cells, and decipher regulators of Th17 effector functions *in vitro* and *in vivo*. We compared the chromatin landscape of non-pathogenic and pathogenic Th17 cells to each other and to other CD4^+^ T cell subsets revealing pTh17 cells as an intermediate between npTh17 and Th1 cells with a substantial overlap with the Th1 regulatory program. We predicted novel drivers of Th17 pathogenicity by analyzing matched scATAC-seq and scRNA-seq and by chromatin and RNA profiling along three polarization time courses. Lastly, we identified and validated the transcription factor BACH2 as a novel suppressor of Th17 pathogenicity *in vitro* and *in vivo* and showed relevant genetic evidence for protective variants in the BACH2 locus associated with MS disease risk in humans and with chromatin organization and higher BACH2 expression in T cells.

Our discovery of distinct chromatin landscapes of npTh17 and pTh17 cells suggests major chromatin differences between them and supports a model in which these represent distinct cell fates. The concept of beneficial and pathogenic Th17 cells is now well established ^2, 4, 81^, associated with expression differences between npTh17 and pTh17 cells ^6, 8, 16, 82^ and different regulators ^29, 33, 48, 63^. However, whether npTh17 and pTh17 cells represent plastic cell states or stable cell fates remained elusive.

The chromatin signature distinguishing npTh17 from pTh17 cells *in vitro* and *in vivo* allowed us to identify mechanisms by which these distinct cellular fates are achieved. Our analysis of the chromatin accessibility in individual npTh17 and pTh17 cells, and along polarization time courses, showed how the two diverge from each other during polarization and allowed us to discover multiple novel putative regulators of Th17 pathogenicity. Several regulators, such as *Bach2* and *Atf3*, were not appreciated in previous studies that only analyzed RNA profiles, highlighting the benefit of chromatin accessibility analysis combined with motif analysis when searching for novel regulators. Most differences in chromatin accessibility between npTh17 and pTh17 cells were identified in intronic and intergenic regions (>92%), demonstrating how distal ACRs are critical components of regulatory programs that drive cell fate. The substantial differences also open up the possibility of targeting pTh17 cells specifically in autoimmune diseases, leaving the beneficial functions of npTh17 cells intact. The chromatin signatures derived here with matched putative target genes by single-cell analysis provide a rich resource to find putative regulatory elements and pathways that can be modified to specifically alter Th17 cell states and tune Th17 cell function.

pTh17 cells have previously been shown to up-regulate Th1-marker genes (e.g. *Tbx21* and *IFNg*)^4, 81^, in both disease models in mice ^8, 16, 18^ and in human disease ^9, 20–22^. However, it remained unclear whether pTh17 cells up-regulate only a few Th1-marker genes or acquire entire Th1-like cellular programs, and if so, what genes and regulators belong to this Th1-like pathogenic Th17 program. Our analysis discovered a major regulatory network, at the chromatin, mRNA and regulator level, that is shared between pTh17 cells and Th1 cells and positions pTh17 cells as an intermediate cell fate between npTh17 cells and Th1 cells. We defined a Th1-like pathogenic Th17 subnetwork and identified novel pathogenic effector and regulator molecules, including BACH2. Our study sets the foundation for future studies aiming to target these novel effector and driver molecules as novel treatment options for autoimmune diseases.

BACH2 is a transcriptional repressor of the basic region leucine zipper (bZIP) TF family ^74^, and was shown to be required for the differentiation and homeostasis of Treg cells by repressing effector cell differentiation ^73, 83^ and in CD8^+^ T cells to play an important role in memory formation after acute infection ^84, 85^. However, the function of BACH2 in Th17 cells has not been studied. Here, we identify BACH2 as a novel repressor of Th17 pathogenicity, such that BACH2 deficiency leads to an induction of the Th1-like pathogenic program in npTh17 cells, whereas BACH2 overexpression suppresses Th17 pathogenicity *in vivo* and converts pathogenic Th17 cells into stem-like Th17 cells that we have recently described ^16^. Interestingly, a recent study demonstrated an important role of BACH2 in the generation of stem-like CD8^+^ T cells ^76^. We and others have previously shown that npTh17 cells share multiple properties with stem-like CD8^+^ T cells, including a shared expression profile ^8, 16, 86^. This supports a model in which BACH2 acts as a mediator of the stem-like, non-pathogenic Th17 program and its overexpression in pTh17 cells suppresses and prevents their ability to shift into a highly pathogenic effector state by impacting the chromatin landscape and expression of the Th1-like program in pTh17 cells.

We found strong support for the relevance of the role of BACH2 in human autoimmune disease. Single-nucleotide polymorphisms (SNPs) at the *BACH2* locus are associated with multiple autoimmune diseases, the protective allele for MS susceptibility results in an increase of *BACH2* expression in CD4^+^ T cells, and the locus contains a prominent T cell super-enhancer ^77, 87, 88^. Moreover, BACH2 has previously been predicted to regulate the transcriptional program of human TH17-IL-10^+^ cells ^89^. Hence, an induction of BACH2 expression and/or activity leading to the generation of homeostatic Th17 cells in human autoimmune diseases might yield a novel, promising treatment approach for inhibiting pathogenic effector T cells.

In conclusion, we have used comprehensive analysis of the chromatin accessibility and associated gene expression of np and pTh17 cells and other CD4^+^ T cell subsets to identify a novel Th1-like pathogenic Th17 program that is shared between pTh17 cells and Th1 cells at both the chromatin and expression levels, and is governed along with other distinct programs by an integrated regulatory network with a self-reinforcing mutually exclusive architecture. We also predicted and validated BACH2 as a novel regulator of Th17 pathogenicity. Our work provides a framework to leverage chromatin profiles to yield novel insights into the regulation of CD4^+^ T cell diversification and yields a foundation for future explorations of novel drivers of Th17 pathogenicity.

## Acknowledgments

We thank the Broad Genomics Platform for help with bulk RNA-seq sample processing, and Leslie Gaffney with graphical help in Fig. preparation, and Mary Collins for critical feedback on the manuscript. We thank all members of the Kuchroo and Regev labs for feedback. We thank Junrong Xia, Helene Stroh, Rajesh Kumar Krishnan, and Deneen Kozoriz for technical support. A.S. was supported by a German Academic Scholarship Foundation (Studienstiftung des Deutschen Volkes) PhD fellowship. P.I.T. was supported by an NIH F32 Ruth L. Kirschstein Postdoctoral Fellowship (5F32AI138458). This work was supported by National Institute of Health grants (R01NS045937, R01NS30843, R01AI144166, P01AI073748, P01AI039671 and P01AI056299) (V.K.K.) and by the Klarman Cell Observatory and HHMI (A.R.). F.S. was supported by the Swiss National Science Foundation (grant n. 189331). F.S. and the Institute for Research in Biomedicine are supported by the Helmut Horten Foundation. N.A.P. was supported in part by National Multiple Sclerosis Society (grants JF-1808-32223 and RG-1707-28657).

## Author contributions

P.I.T., A.S., V.K.K. and A.R. conceived the study. P.I.T., A.S., O.R.R., V.K.K. and A.R. designed the experiments and interpreted the results. P.I.T and A.S. performed the *in vitro* T cell culture experiments. A.S. performed with assistance from Y.H. the *in vivo* mouse experiments. P.I.T and A.S. performed the ATAC-seq and RNA-seq experiments, with assistance from M.Z., E.C., S.Z., C.W., Va.S., Ve.S, and O.R.R.. P.I.T. designed and performed the computational analysis and L.H. performed the network analysis with assistance and guidance from A.S., V.K.K., and A.R.. S.M. and J.D.B. helped with scATAC-seq experiments. S.N. and F.S. helped with the acquisition of human T cell samples. N.A.P. provided the human *BACH2* GWAS analysis. The manuscript was written by A.S. and P.I.T. and was edited by V.K.K. and A.R. with input from all authors.

## Table legends

Table S1: Bulk ATAC-seq and RNA-seq signal from gene-to-peak correlations in npTh17 vs. pTh17 cells

Table S2: Bulk ATAC-seq of CNS-derived *vs.* dLN-derived Th17 cells

Table S3: Bulk RNA-seq of CNS-derived *vs.* dLN-derived Th17 cells

Table S4: GSEA for genes near differentially accessible peaks in cells harvested from the CNS *vs.* dLN during EAE

Table S5: Bulk ATAC-seq of CD4^+^ T cell subsets

Table S6: Bulk RNA-seq of CD4^+^ T cell subsets

Table S7: Bulk ATAC-seq time course of Th17 differentiation

Table S8: Peak-to-gene links of scATAC/RNA-seq of Th17 and Th1 cells

Table S9: TF regulators from integrated scATAC-seq and scRNA-seq of Th17 and Th1 cells

Table S10: Bulk ATAC-seq of *Bach2* KO Th17 and Th1 cells

Table S11: GSEA for genes near differentially accessible peaks in *Bach2* KO *vs.* control Th17 cells

Table S12: Bulk ATAC-seq of *Bach2* OE Th17 and Th1 cells

Table S13: GSEA differentially expressed genes in *Bach2* OE *vs.* control cells harvested from the CNS during EAE

## Methods

### Mice

C57BL/6J wild-type, C57BL/6-*Il17a^tm1Bcgen^*/J (named *Il17a^GFP^*) mice, B6.SJL-*Ptprc*^a^*Pepc*^b^/BoyJ (named CD45.1), and B6(C)-*Gt(ROSA)26Sor*^em1.1(CAG-cas9*,-EGFP)Rsky^/J (named Cas9 knockin mice) were obtained from the Jackson Laboratory. C57BL/6-Tg(Tcra2D2,Tcrb2D2)1Kuch/J (named 2D2) mice were generated in our laboratory ^75^. All mice were housed under specific-pathogen-free (SPF) conditions at the Brigham and Women’s Hospital (BWH) mouse facility in Boston. All experiments were carried out in accordance with and approved by the guidelines of the Brigham and Women’s Hospital (BWH) Institutional Animal Care and Use Committee (IACUC) in Boston.

### Primary cell culture

Primary cells were cultured at 37°C and 10% CO_2_ in a humidified incubator in complete medium that consisted of DMEM medium (Gibco, Cat# 11-965-118) supplemented with 10% fetal bovine serum, sodium pyruvate (Gibco), L-glutamine (Gibco), penicillin/streptomycin (Gibco), MEM non-essential amino acids solution (Gibco), L-arginine/L-asparagine (Sigma Aldrich), folic acid (Sigma Aldrich), MEM vitamin solution (Gibco), 2-mercaptoethanol (Sigma Aldrich), and gentamicin sulfate (Lonza BioWhittaker).

### Isolation and differentiation of naïve CD4^+^ T cells

CD4^+^ T cells were isolated from the spleen and peripheral lymph nodes using anti-CD4 microbeads (Miltenyi Biotec) following the manufacture’s protocol. Naïve CD4^+^ CD44^-^ CD62L^+^ cells were sorted using a BD FACS Aria IIIu flow cytometer (BD Biosciences). Naïve CD4^+^ T cells were activated with plate-bound anti-CD3 (1μg/ml, clone 145-2C11, Bio X Cell) and anti-CD28 (1μg/ml, clone PV-1, Bio X Cell) antibodies on flat-bottom 96-well plates in the presence of cytokines. Non-pathogenic Th17 cell differentiation was performed with IL-6 (25 ng/mL, R&D Systems) and TGF-β1 (2 ng/mL, Sigma Aldrich). Pathogenic Th17 cells were differentiated with IL-1β (20 ng/mL, R&D Systems), IL-6 (25ng/mL) and IL-23 (20ng/mL, R&D Systems). For Th1 differentiation cultures, the medium was supplemented with IL-12 (20ng/mL, R&D Systems). Differentiation of T_reg_ cells was performed with TGF-β1 (2ng/mL).

### Active induction of experimental autoimmune encephalomyelitis (EAE)

6- to 8-week-old sex-matched mice were immunized subcutaneously into the flanks with an emulsion of the MOG35-55 peptide (100μg/mouse, Genemed Synthesis) and *M. tuberculosis* H37Ra extract (5mg/ml, Becton Dickinson) in CFA (200ul/mouse, Becton Dickinson). Pertussis toxin (100ng/mouse, List Biological Laboratories) was injected intravenously on day 0 and day 2 post immunization. EAE disease course was monitored daily, and each mouse was assigned a clinical score according to the following criteria: 0, no symptoms; 1, limp tail; 2, impairment in righting reflex and/or hind limb weakness; 3, hind limb paralysis; 4, total limb paralysis; 5, moribund or dead. All mice presenting a clinical score >4 were euthanized.

### Retroviral transduction in primary CD4^+^ T cells

For the retrovirus production, HEK293T cells were transfected with gag/pol (1 μg/ml) and pCL-Eco (0.5 ng/ml) retroviral plasmids and the transfer plasmids containing *Bach2*-targeting gRNA or non-targeting gRNA (retro-gRNA-mRFP1, 1.3 μg/ml, marked by dsRed), and *Bach2* ORF (Origene, MR224703 cloned into Addgene, Cat# 52107, 1 μg/ml, marked by GFP) using the X-tremeGENE transfection reagent (Sigma Aldrich) following the manufacturer’s directions. Media was changed 12-14hr after transfection, and 36hr later, cell culture supernatant was collected and used for transduction of primary CD4^+^ T cells. T cells were activated with plate-bound anti-CD3 and anti-CD28 antibodies (1μg/ml) in complete media in the presence of polarization cytokines and transduced at 24h after activation by spin transduction with polybrene (Santa Cruz Biotechnology, 8 μg/ml) at 2000 rpm, for 2hr at 35°C. To confirm CRISPR/Cas9-mediated *Bach2* knockout, genomic DNA was collected from treated cells and the region surrounding the gRNA target site was amplified. Indel efficiency was measured by Sanger sequencing trace decomposition, comparing targeting gRNA-treated samples to non-treated ^90^. To validate *Bach2* overexpression by qPCR, total RNA was extracted using the PicoPure RNA Isolation Kit (Applied Biosystems) and cDNA was synthesized with the SuperScript VILO cDNA Synthesis Kit (Invitrogen). QPCR was conducted using the TaqMan Fast Advanced Master Mix (Applied Biosystems) in the ViiA 7 Real-Time PCR system (Applied Biosystems). The following TaqMan probes (ThermoFisher Scientific) were used: *Bach2* (Cat# Mm00464379_m1), *18S* (Cat# 4352930E), and *Actb* (Cat# 4352341E). Relative expression levels are depicted as 2^-ΔCT^ values, ΔCT = (gene of interest CT) - (geoMean Housekeeper CT).

### Adoptive transfer EAE

Naïve 2D2 transgenic T cells were differentiated *in vitro* for 7 days. Cells were first activated with plate-bound anti-CD3 and anti-CD28 antibodies (1μg/ml) in complete media containing differentiation cytokines for the pTh17 condition, rested for two days, and then reactivated for two days with anti-CD3 and anti-CD28 antibodies (1μg/ml) in complete media without cytokines. For the transfer of *Bach2* OE cells, cells were transduced as described above. 2×10^6^ cells were transferred intravenously into congenically marked, wild-type (CD45.1) recipients.

### Isolation of lymphocytes from EAE mice

Mice were sacrificed at the peak of EAE disease. Perfusion was performed intracardially with cold PBS. Brain and the spinal cord were flushed out with PBS by hydrostatic pressure. CNS was minced with a razor blade and digested with collagenase D (2.5mg/ml, Roche Diagnostics) at 37°C for 20 min. After passing the tissue through a 40 μm strainer, mononuclear cells were enriched by centrifugation through a Percoll gradient (37% and 70%). Mononuclear cells were harvested from the interphase and used for downstream analysis. Cells from draining lymph nodes were obtained by mashing lymph nodes through a 40 μm strainer.

### Flow cytometry and fluorescence-activated cell sorting (FACS)

For flow cytometric analysis and FACS, single-cell suspensions were stained in flow buffer (2% fetal bovine serum in PBS) with corresponding antibodies for surface proteins for 30min at 4°C in the dark. Antibodies with specificity for the following cell surface proteins were purchased from Biolegend with different fluorochrome labels: CD4 (clone RM4-5), CD25 (clone PC61), CD44 (clone IM7), CD62L (clone MEL-14), TCRβ (clone H57-597). Viability staining was performed using the eF506 dye (eBioscience). Intracellular cytokine staining was performed by activating cells for 4-5h at 37°C with the Cell Stimulation Cocktail (plus protein transport inhibitors) (eBioscience) and subsequent fixing and staining per manufacturer’s instructions using the BD Fixation/Permeabilization Solution Kit (BD Biosciences). For intranuclear transcription factor staining, the FoxP3/Transcription Factor Staining Buffer Set (eBioscience) was used. Intracellular and intranuclear antibody staining was performed for 30min at 4°C in the dark. Antibodies with specificity for the following intracellular and intranuclear proteins were purchased from Biolegend with different fluorochrome labels: GM-CSF (clone MP1-22E9), IFNγ (clone XMG1.2), IL-10 (clone JES5-16E3) IL-17A (clone TC11-18H10.1). For staining of FoxP3, clone FJK-16s was purchased from eBioscience. For flow cytometry analysis, samples were acquired on BD LSRII (BD Biosciences) or BD LSRFortessa (BD Biosciences) flow cytometers. FACS was performed on a BD FACS Aria IIIu flow cytometer (BD Biosciences). Flow cytometry data were analyzed with FlowJo software (FlowJo LCC) and GraphPad Prism software (GraphPad).

### Bulk ATAC-seq

6,000 viable cells were sorted into PBS/2% FCS and were stored in Bambanker^TM^ cell freezing media (LYMPHOTEC Inc) at -80°C. For library preparation, cells were thawed at 37°C, washed with PBS, and lysed and tagmented in 40μl of 1X TD Buffer, 0.2μl TDE1 (Illumina), 0.01% digitonin, and 0.3X PBS as previously described ^25^. Transposition was performed at 300 rpm and 37°C for 30min followed by DNA purification with the MinElute PCR Purification Kit (QIAGEN). Eluate was immediately amplified by PCR. In the first step, 5 cycles of pre-amplification were performed using NEBNext High-Fidelity 2X PCR Master Mix (NEB) with indexed primers (Ad1 and Ad2). Additional cycle numbers were determined by SYBR Green (Invitrogen) qPCR. Libraries were then purified with the MinElute PCR Purification Kit (QIAGEN) and library quantification was performed with the KAPA Library Quantification Kit (KAPA Biosystems) and a Qubit dsDNA HS Assay kit (Invitrogen). Final libraries were sequenced on an Illumina NextSeq 550 system with paired-end reads of 37 base pairs in length.

### Bulk RNA-seq

6,000 cells were sorted by FACS into 5μl TCL buffer (QIAGEN) supplemented with 1% 2-mercaptoethanol (Sigma Aldrich). Libraries were prepared using a modified SMART-Seq2 protocol ^91^, including RNA secondary structure denaturation at 72°C for 3mins, reverse transcription with Maxima Reverse Transcriptase (ThermoFischer Scientific), and whole transcriptome amplification (WTA) using the KAPA HiFi HotStart ReadyMix 2X (KAPA Biosystems) for 12 cycles. WTA product quality was confirmed with a High Sensitivity DNA Chip run on a Bioanalyzer 2100 (Agilent, Cat# G2939BA) and quantified using a Qubit dsDNA HS Assay Kit (Thermo scientific). Libraries were prepared with a Nextera XT DNA Library Preparation Kit (Illumina) and sequenced using a high output V2 75 cycle kit (Illumina) on a NextSeq 500 sequencer (Illumina) with 2×38 paired-end reads.

### Single-cell ATAC-seq

Naïve CD4^+^ T cells were *in vitro* differentiated to npTh17, pTh17, and Th1 cells as described above. After 48h, live cells were sorted into PBS/2%FCS. Cells were lysed in 10 mM Tris-HCl pH 7.4, 10 mM NaCl, 3 mM MgCl2, 0.1% NP40, 0.1% Tween20, 0.01% Digitonin, 1% BSA and permeabilized for 3 min on ice. Permeabilized cells underwent tagmentation and were encapsulated into droplets and libraries were prepared using the Chromium Single Cell ATAC Library & Gel Bead Kit (10x Genomics) following the manufacturer’s instructions. Each *in vitro* differentiation condition was separately loaded onto a 10X channel with n = 2 replicates for each condition. Libraries were sequenced on a NextSeq 500 sequencer (Illumina) with 2 x 50 paired-end reads.

### Single-cell RNA-seq

Naïve CD4^+^ T cells were *in vitro* differentiated to npTh17, pTh17, and Th1 cells as described above. After 48h, live cells were sorted into PBS/2%FCS and were encapsulated into droplets and libraries were prepared using the Chromium Single Cell 5’ Library & Gel Bead Kit (10x Genomics) following the manufacturer’s instructions. Each *in vitro* differentiation condition was separately loaded onto a 10X channel with n = 2 replicates for each condition. Libraries were sequenced on a HiSeq X (Illumina) with paired-end reads of 28 cycles for read 1 and 91 cycles for read 2.

#### Computational methods

##### Bulk ATAC-seq pre-processing

Reads were aligned, filtered for duplicates and mitochondrial reads, and assessed for quality control metrics using the publicly available ENCODE ATAC-seq pipeline (https://github.com/ENCODE-DCC/atac-seq-pipeline version 1.5.4). Briefly, fastq files were trimmed based on detected adapter sequences by cutadapt -m 5 -e 0.2. Bowtie2 was used to align reads to the mm10 reference genome (bowtie2 -X2000 -mm) ^92^. Resulting BAM files were filtered to remove PCR duplicates using PicardTools and to remove mitochondrial reads. Filtered BAM files were merged by condition and peaks were called using MACS2 (macs2 callpeak –nomodel – nolambda –keep-dup all –call-summits) ^93^. A counts table of reads per peak was generated using the bedtools multicov tool ^94^. Read counts were normalized using DESeq2 ^95^. Data were transformed using regularized log transformation with DESeq2 prior to visualization with principal component analysis (PCA). PCA was performed on the regularized log transformation (function: rld) of peak counts from DESeq2 using the prcomp function in R and the top 50,000 peaks. Differential peak analysis between conditions was performed by modeling the count data as a negative binomial generalized linear model and using Wald’s test for significance followed by Benjamini-Hochberg FDR. Differentially accessible peaks were defined as FDR <0.05 and log_2_(fold change)| >0.5. For heatmaps, data were visualized as a z-score across all conditions of log-normalized counts from DESeq2, unless otherwise noted. *K*-means clustering was performed using the kmeans function in R and number of clusters was determined empirically and is specified in the relevant Fig.s. Peak annotation with target genes was performed using GREAT (version 4.0.4, http://great.stanford.edu/public/html/) with default parameters ^34^. Motif enrichment analysis was performed on bed files from selected differentially accessible peaks using homer findMotifsGenome v4.10 (http://homer.ucsd.edu/homer/motif/), with a bed file of peaks as input, -size given.

##### Bulk RNA-seq pre-processing

Reads were aligned to the mm10 genome using Tophat (v2.1.1, https://ccb.jhu.edu/software/tophat/index.shtml) ^96^ and after duplicate removal, counts tables of genes by sample were generated by htseq counts. Read counts were normalized using DESeq2 ^95^. Data were transformed using regularized log transformation with DESeq2 prior to visualization with principal component analysis (PCA). PCA was performed on the regularized log transformation (function: rld) of gene counts from DESeq2 using the prcomp function in R and the top 5,000 genes. Differential gene expression analysis was performed by modeling the count data as negative binomial generalized linear model and using Wald’s test to test for significance followed by Benjamini-Hochberg FDR. Differentially accessible genes were defined as FDR <0.05 and |log_2_(fold change)| >0.5, unless otherwise noted. For heatmaps, data were visualized as a z-score across all conditions of log-normalized counts from DESeq2, unless otherwise noted.

##### Bulk ATAC-seq peak visualization

To generate ATAC-seq signal tracks, fold enrichment bedgraph files were generated from merged bam files per condition using macs2 callpeak (-B -SPMR) followed by macs2 bdgcmp (-m FE). Resulting bedgraph files were sorted and converted to bigWigs using bedGraphToBigWig (https://www.encodeproject.org/software/bedgraphtobigwig/). Tracks were visualized using pyGenomeTracks with a defined genome interval (https://github.com/deeptools/pyGenomeTracks). To generate heatmaps and line profiles of ATAC-seq peaks, bigwig files were converted to a matrix using deeptools’ (v3.4.3, https://deeptools.readthedocs.io/en/develop/) computeMatrix (--referencePoint center -b 1000 -a 1000) with a bed file of selected peaks as input. Deeptools plotHeatmap (--perGroup) and plotProfile (--perGroup) were used to generate heatmaps and line profiles respectively.

##### Gene signature scoring on population ATAC-seq

When scoring genes based on ATAC-seq signal, differentially accessible peaks were assigned to genes by GREAT as described above. Signature scoring was performed with the fgsea package in R using the DESeq2 with a Wald test statistic and nperm = 10000. Each signature was split into a positive module of upregulated genes and a negative module of downregulated genes, and the overall signature score was calculated as the difference between the positive module signature score and the negative module signature score.

##### Gene-to-peak correlation for bulk ATAC-seq and RNA-seq

Gene-to-peak correlation was calculated for the top differentially expressed genes between npTh17 and pTh17 conditions at 24h, 48h, and 72h. Top differentially expressed genes were selected for each time point separately and defined as the 100 genes with highest log_2_(fold change) and 100 genes with lowest log_2_(fold change) when comparing pTh17 to npTh17 cells in the same time point (FDR <0.05). The union was taken between the three time points. For each differentially expressed gene, a gene window was defined as +/- 100kb around the gene transcription start site (TSS). These genes were compared with the top differentially accessible peaks selected at each time point and defined as the 100 peaks with highest log_2_(fold change) and 100 peaks with lowest log_2_(fold change) when comparing pTh17 and npTh17 cells at the same time point (FDR <0.05). All differentially accessible peaks that intersected this gene window by at least one base were initially assigned to the gene. The Pearson correlation coefficent between normalized RNA-seq read counts and normalized ATAC-seq read count nearby differentially accessible peaks was calculated, and the differentially accessible peak within the 200kb window that had the highest correlation was matched to the gene. As a null model, the same number of non-differentially expressed genes from similar expression bins were randomly selected and randomly matched to non-differentially accessible peaks of similar accessibility bins and correlations were calculated. For the null model, data were partitioned into 10 bins based on average values across all conditions. Only peak-to-gene matches with positive correlations (r >0) with a Benjamini-Hochberg FDR <0.05 in a t-test comparing to the null model were selected. Data were visualized as a z-score across all conditions of log-normalized counts from DESeq2 (described in Bulk ATAC pre-processing).

##### Comparison of bulk ATAC-seq and RNA-seq of *in vivo*-derived Th17 cells

Bulk ATAC-seq and RNA-seq from CNS- and dLN-derived Th17 cells were pre-processed and visualized in tracks as described above. To relate the *in vivo* chromatin accessibility changes to those observed *in vitro*, all peaks called from the *in vivo* ATAC-seq dataset (dLN and CNS) were intersected with all peaks from the *in vitro* pTh17 and npTh17 ATAC-seq dataset (at time points 24h, 48h, and 72h) using bedtools (v2.26.0) intersectBed (-wa -f 0.75) to determine peaks with a minimum 75% overlap. From this shared peak-space, peaks were identified that were significantly higher both in CNS *vs*. dLN Th17 cells (FDR <0.05) and in pTh17 *vs*. npTh17 cells (FDR <0.05 using the 48h time point), or that were significantly higher both in dLN *vs* CNS Th17 cells (FDR <0.05) and in npTh17 *vs* pTh17 cells (FDR <0.05).

##### Time course analysis of *in vitro* CD4 T cell polarization

Population ATAC-seq and RNA-seq from CNS- and dLN-derived Th17 cells were pre-processed as described above. Specifically, for ATAC-seq, peaks were called from merged bam files from each condition at each time point and the union of these peak sets was defined as the universe of peaks to generate counts tables. Chromatin accessibility was compared for each pair of cell types (npTh17 and pTh17; pTh17 and Th1; npTh17 and Th1) at each time point. The top 500 changes were selected from each comparison (FDR <0.05, ranked by log_2_(fold-change)) and the union of these sets was defined as the differentially accessible peak set. To prioritize shared chromatin accessibility changes between any two groups in the dataset, peaks were identified that were differentially more open in each of any two cell subsets compared to the third at a given time point (*e.g.*, peaks that were more accessible in both npTh17 and pTh17 cells compared to Th1 cells). The union of the top 500 of each comparison (FDR <0.05, ranked by average log_2_(fold-change)) was defined as a shared peak-set. The pair-wise differentially accessible peak set and shared-peak-set were combined and z-score of log-normalized counts were plotted for heatmap visualization. Clustering was performed using kmeans in R (k=10, determined empirically). To relate differentially accessible peaks from clusters to the principal components (PCs) related to time, activation, and polarization condition, peaks were assigned to PC1, PC2, and PC3 if their absolute loadings were in the top 20% of loadings. If a peak’s loadings were in the top 20% of multiple PCs, it was assigned to the PC with maximal absolute loading. This peak-set was intersected with the peaks assigned to each cluster to assign a PC (PC1, PC2, PC3, or none).

##### Single-cell RNA-seq data pre-processing

ScRNA-seq pre-processing was performed using the Cumulus analysis workflow version 0.11.0 using CellRanger v2.2.0 ^97^. Raw sequencing data were demultiplexed using CellRanger mkfastq and aligned to the reference mm10 (v 1.2.0) and a gene counts table was generated using CellRanger count (expect-cells = 3000). Cell by counts tables were combined for six channels (n = 2 of npTh17, pTh17, and Th1 cells) using CellRanger aggr for downstream analysis.

##### Single-cell RNA-seq data analysis

Downstream analysis, including normalization and clustering, was performed using Seurat v3.1.5, as follows ^98^. The gene counts matrix was used as input to Seurat and low-quality cells and doublets were filtered by number of genes >1000 and <6000, mitochondrial reads <3%, and number of unique molecules detected >3000. Gene counts were normalized by total expression, scaled by 10000, and log transformed. Next, data were scaled using ScaleData. Principal component analysis was performed on the top 2000 variable features identified by FindVariableFeatures (selection.method = “vst”). A *k*-nearest-neighbor (*k*-NN) graph was generated with *k*=20 and the top 10 principal components, clustered with the Leiden algorithm using FindClusters (resolution 0.5), and embedded using RunUMAP.

##### Single-cell ATAC-seq pre-processing

ScATAC-seq pre-processing was performed using the Cumulus analysis workflow version 0.11.0 running CellRanger-ATAC v1.0.1 ^97^. Raw sequencing data were demultiplexed using CellRanger-ATAC mkfastq and aligned to the reference mm10 genome (mm10_atac_v1.0.1). Cell by counts tables were combined for six channels (n = 2 of npTh17, pTh17, and Th1 cells) using CellRanger aggr for downstream analysis.

##### Single-cell ATAC-seq data analysis

Downstream analysis, including clustering, gene scoring, integration with matched scRNA-seq data, DNA peak-to-gene linkage, and transcription factor motif deviation scoring, was performed using ArchR v1.0.1 ^57^ (according to manual instructions, https://www.archrproject.com/bookdown/index.html#section) and Seurat v3.1.5 ^98^, as follows. Low quality cells were filtered based on number of unique peak fragments (nFrags >1500 and nFrags <75000) and signal-to-noise ratio (transcription start site enrichment >5), retaining a total of 27,515 cells. A gene score matrix was generated by summing accessibility signal within the 100kb window on either side of each gene. Dimensionality reduction was performed using ArchR’s implementation of latent semantic indexing, addIterativeLSI (iterations = 2, resolution = 0.5, varFeatures = 25000) and data were visualized by addUMAP (nNeighbors = 30). Of eight clusters identified using this method, two (“mixed” and “npTh17_2”) were removed from further analysis based on abnormally high fragments detected, suggesting doublets or low-quality cells. Cluster markers were identified using getMarkerFeatures (FDR <=0.05, log_2_(fold change) >0.2, testMethod = “wilcoxon”). For visualization of sparse ATAC signal on UMAP, gene scores were imputed by addImputeWeights. Peaks were called on each cluster and on cells grouped by treatment condition using addGroupCoverages, which is a wrapper of macs2. Gene-set signature scores were calculated using addModuleScore, a wrapper from Seurat ^98^, with the gene score matrix as input.

##### Integrating matched scATAC-seq and scRNA-seq datasets for peak-to-gene linkage

Single cell profiles in the scATAC-seq data set were aligned to profiles from the scRNA-seq dataset by ArchR’s integration method ^57^. Briefly, a gene integration matrix is generated by addGeneIntegrationMatrix, a function based on Seurat’s integration method FindTransferAnchors ^98^, which aligns gene score matrix values from scATAC-Seq data with gene expression values from scRNA-seq data to match each cell from the scATAC-seq dataset with a corresponding scRNA-seq expression profile. Gene integration was constrained to match scRNA-seq profiles to scATAC-seq profiles from cells that were treated with the same polarization condition. This gene integration is used as basis for peak-to-gene linkage analysis for correlation between peak accessibility and gene expression in single cells (addPeak2GeneLinks, Pearson correlation >0.45 and resolution = 1 base-pair).

##### Topic modeling on scATAC-seq data-set

To model cell programs, a Latent Dirichlet Allocation (LDA) model was learned from the scATAC-seq data set using cisTopic (v0.3.0, https://github.com/aertslab/cisTopic) ^58^. Region distributions per topic and topic contributions per cell were modeled using runWarpLDAModels (iterations = 500) using a range for number of topics from 5 to 40. The final number of topics (n=35) was selected by finding the model with the lowest perplexity as a measure of how well the model predicts the data. Topic scores per cell were visualized on the UMAP embedding of the scATAC-seq dataset (as in **Fig. 5A**). The top 500 genome regions representing each topic were identified using binarizecisTopics (method = “Predefined”, cutoffs = 500). Annotation of genomic regions from each topic with target genes was performed using GREAT (version 4.0.4, http://great.stanford.edu/public/html/) with default parameters ^34^.

##### Prediction of transcriptional regulators from single-cell profiling data

Marker peaks for each cell type and cluster (FDR <0.1 and log_2_(fold change) >0.2) were annotated with enriched TF motifs from the Homer (v4.10) database and cisBP with addMotifAnnotations from ArchR (motifSet = “homer”, motifSet = “cisdb”). Motif enrichment was performed using peakAnnotationEnrichment (with FDR <=0.1 and log2(fold-change) >=0.2 set as cut-off parameters), which calculates enrichment of the motif using marker peaks of the test condition compared to background peaks. Adjusted p-values for enrichment were calculated using a hypergeometric test with Bonferroni correction. Motif deviations, a measure of how much the accessibility of peaks with a given motif differs from expected accessibility modeled on a set of background peaks, were calculated using addMotifDeviations, a wrapper of chromVAR ^59^. To find TF regulators of chromatin accessibility, the correlation between motif deviation and gene expression for TFs was found using correlateMatrices. The maximum motif deviation per cluster was plotted by correlation, and TFs with maximum motif delta >2 and correlation >0.4 were selected as candidate positive TF regulators.

##### Regulatory network analysis from scATAC-seq and scRNA-seq

ScATAC-seq and scRNA-seq data were integrated to construct the regulatory network. First, candidate target genes were identified from pairwise differential expression using glmQLFit and glmQLFTest functions in edgeR package ^99, 100^ (FDR <0.01, fold change >2). Next, peaks were linked to candidate target genes using ArchR ^57^ (see **Integrating matched scATAC-seq and scRNA-seq datasets for peak-to-gene linkage**). Peaks were scanned for TF motif binding sites using ArchR (see **Prediction of transcriptional regulators from single-cell profiling data)**. TFs were then associated to each candidate target gene if their binding sites were observed in targets’ linked peaks, based on the homer motif database (v4.10, http://homer.ucsd.edu/homer/motif/) and cisBP database. To refine the network, Pearson correlations of scRNA-seq expression levels of TFs’ transcript and of associated targets were computed across all cells and the connections with low or insignificant correlations were removed ^101^ (restricted to |Pearson correlation| >0.1, FDR <0.001). Finally, the network was further pruned for by restricting to the TFs that regulates at least 10 targets and targets that were regulated by at least 5 TFs. TF and target modules were identified by hierarchical clustering of the rows and columns of a TF-by-target expression correlation coefficient matrix with a Euclidean distance and the Ward1 agglomeration method ^102^. Clustering of targets was restricted to genes that were not detected to function as TFs in the network.

## Supplementary Figure legends

**Fig. S1:**
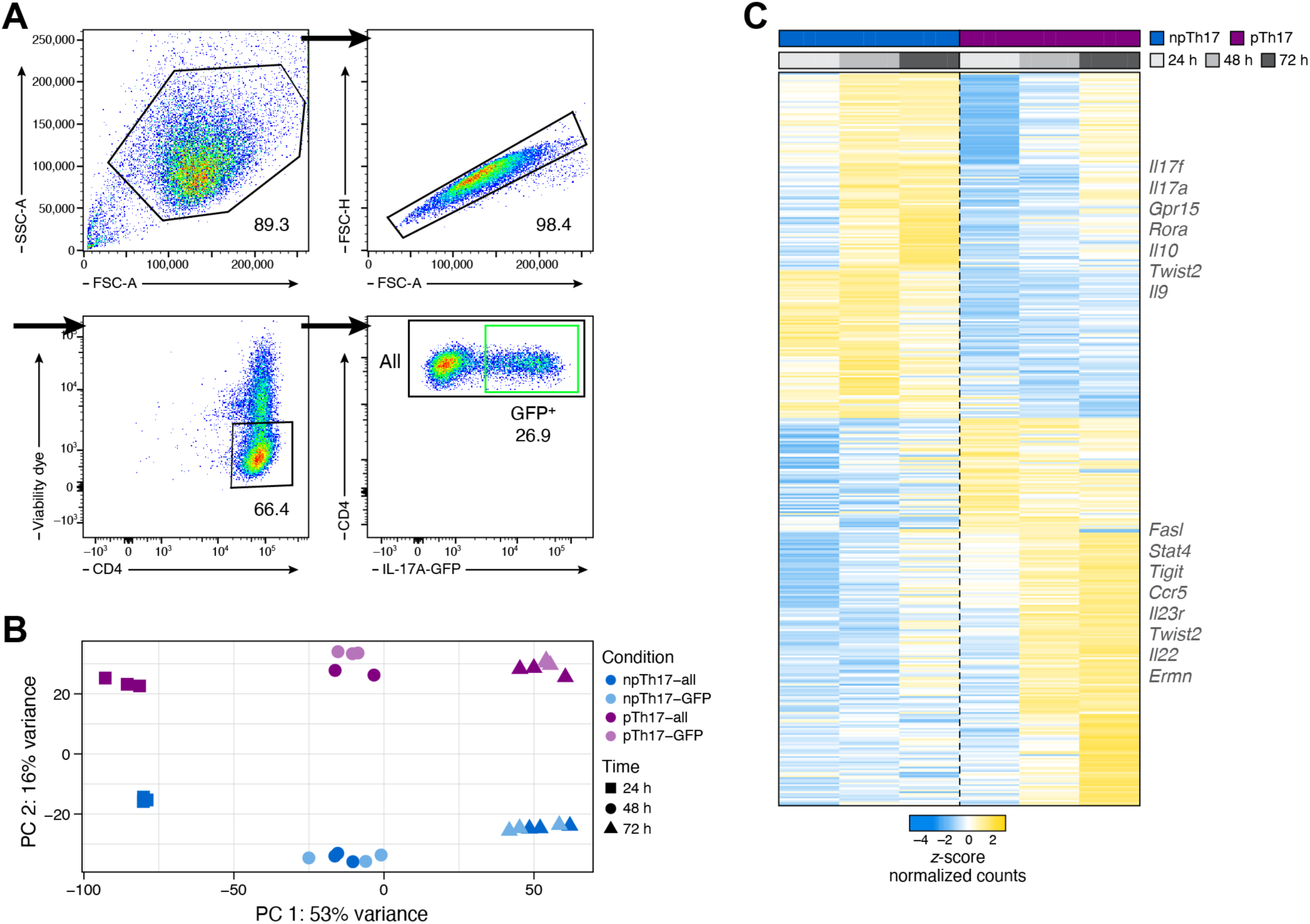
Bulk RNA-seq of *in vitro* differentiated Th17 cells. (**A**) Gating strategy for FACS. Exemplary plot of *in vitro* differentiated npTh17 cells at 72h sorting singlet viable CD4^+^ cells (all, black) and singlet viable CD4^+^ IL-17A-GFP^+^ cells (GFP^+^, green). (**B,C**) Bulk RNA-seq of *in vitro* differentiated npTh17 and pTh17 cell populations. PCA (**B**) of cell profiles and expression (**C,** z-score of normalized counts, color bar) of the union of the top 200 differentially expressed genes (**C**, rows) at 24h, 48h, or 72h (FDR <0.05, rank ordered in C by fold change, 415 genes) in npTh17 (blue) and pTh17 (blue) cells populations from *Il17a*^GFP^ reporter mice either after sorting for GFP^+^ cells (light color) or without sorting (all viable cells, dark color).

**Fig. S2:**
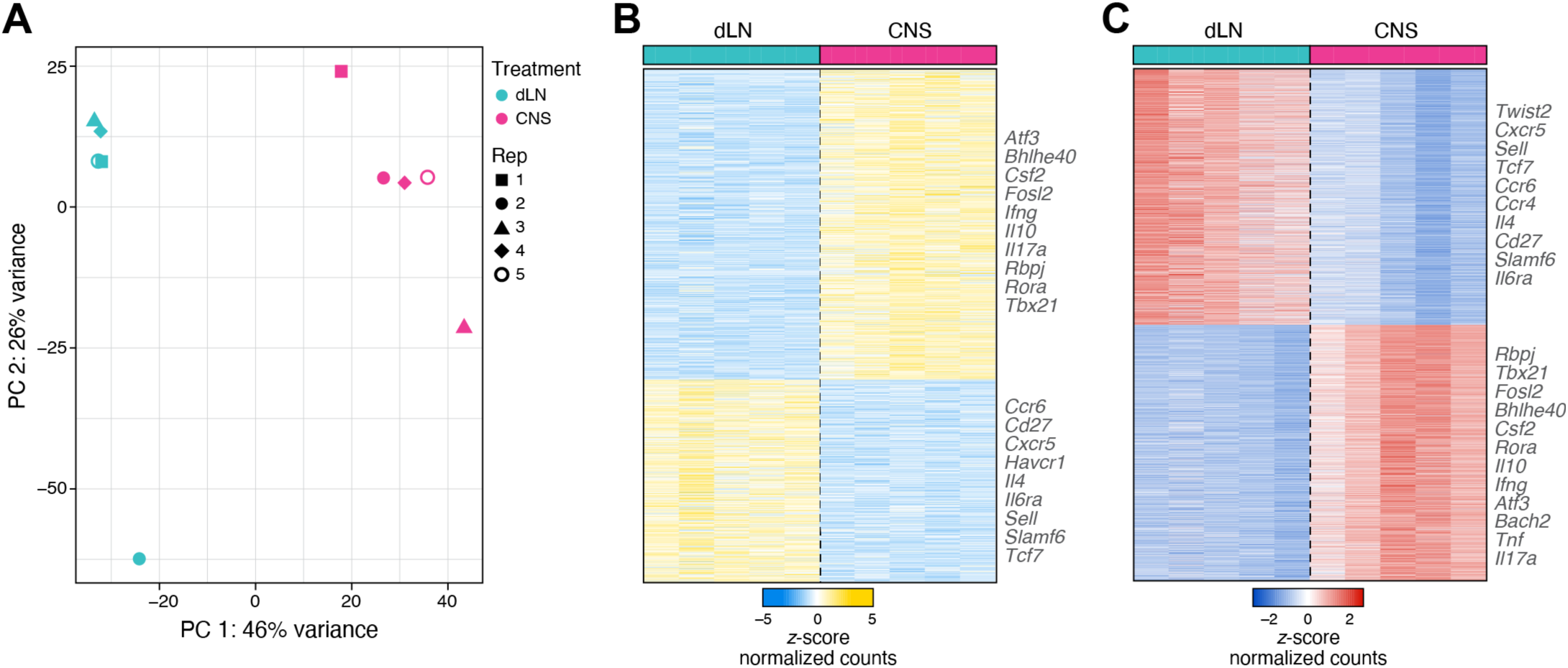
Comparison of *in vitro* and *in vivo* Th17 cell ATAC-seq peaks. (**A,B**) Distinct gene expression profiles of CNS-infiltrating Th17 cells. (**A**) PCA of bulk RNA-seq profiles from viable CD45^+^ TCRβ^+^ CD4^+^ IL17-A-GFP^+^ dLN-derived (teal) and CNS-derived (pink) Th17 cell populations. DLN and CNS samples are matched for each biological replicate (shapes, n=5). (**B**) Relative expression (z-score, color bar) of the 1,565 genes (rows) differentially expressed (FDR <0.05, |log_2_(fold change|) between dLN- (teal) and CNS- (pink) derived Th17 cells (rows). (**C**) Distinct chromatin accessibility profiles of CNS-infiltrating Th17 cells. Chromatin accessibility (z-score of normalized counts, rows) in the top 5,000 DACRs (rows, FDR <0.05) between dLN- (teal) and CNS- (pink) derived Th17 cell populations, ranked by ranked by log_2_(fold change).

**Fig. S3:**
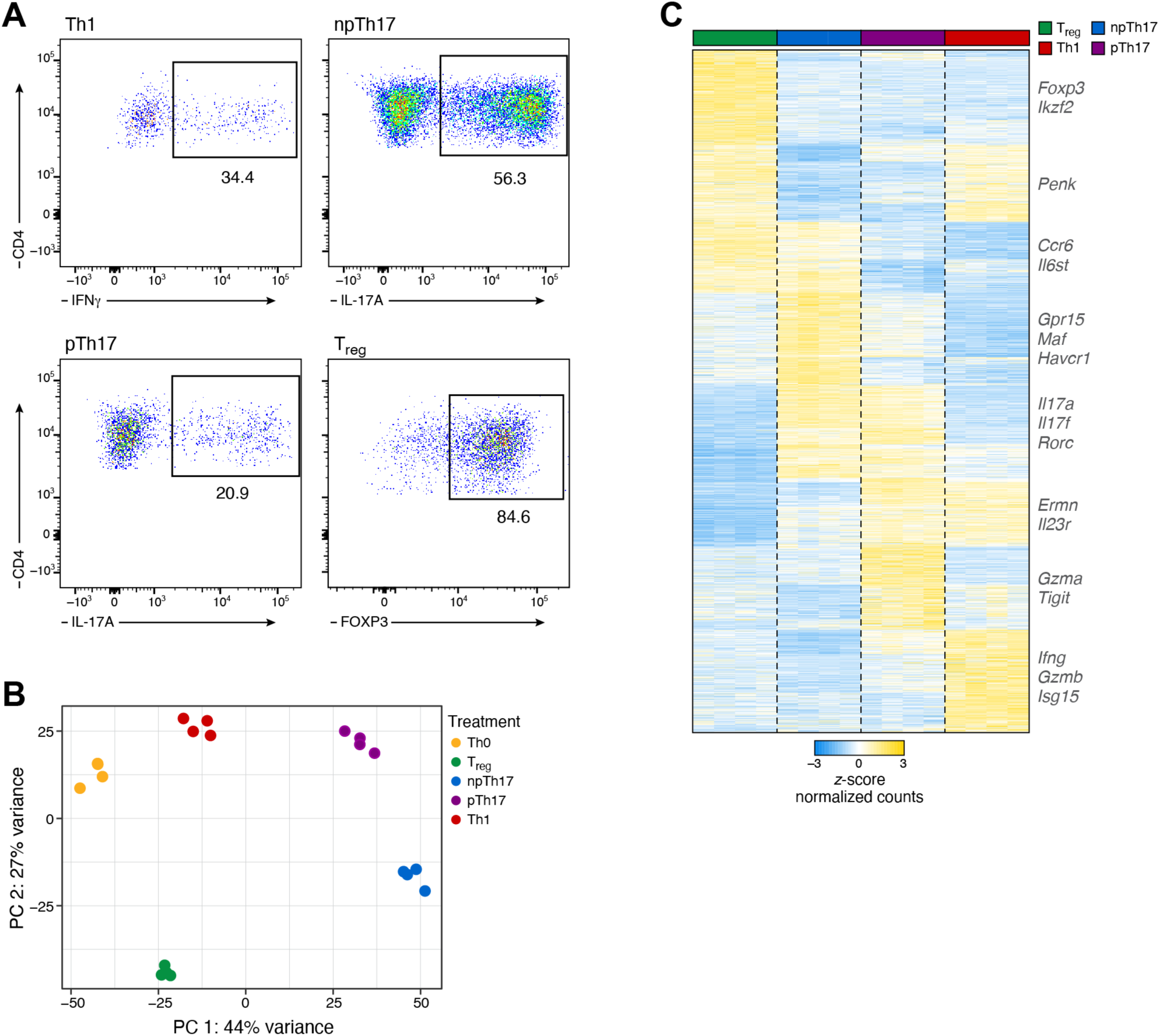
Chromatin landscape of CD4^+^ T cell subsets. (**A**) Expression of signature cytokines in *in vitro* differentiated Th1 cells, npTh17 cells, pTh17 cells, and T_reg_ cells. Illustrative flow cytometry plots are shown. (**B,C**) Distinct gene expression profiles of CD4^+^ T cell subsets. (**B**) PCA (first two PCs) of RNA-seq profiles of *in vitro* differentiated Th0 cells (yellow), npTh17 cells (blue), pTh17 cells (purple), Th1 cells (red), and T_reg_ cells (green) at 72h. (**C**) Differentially expressed genes between pairs of CD4^+^ T cell subsets. Relative expression (z-score) of the top 500 genes (rows) differentially expressed (FDR <0.05) between any two cell-type conditions (1,556 genes total) in cells of each type (columns), with genes ordered by *k*-mean clustering (*k*=8) into cell type specific clusters. Key genes are highlighted on the right.

**Fig. S4:**
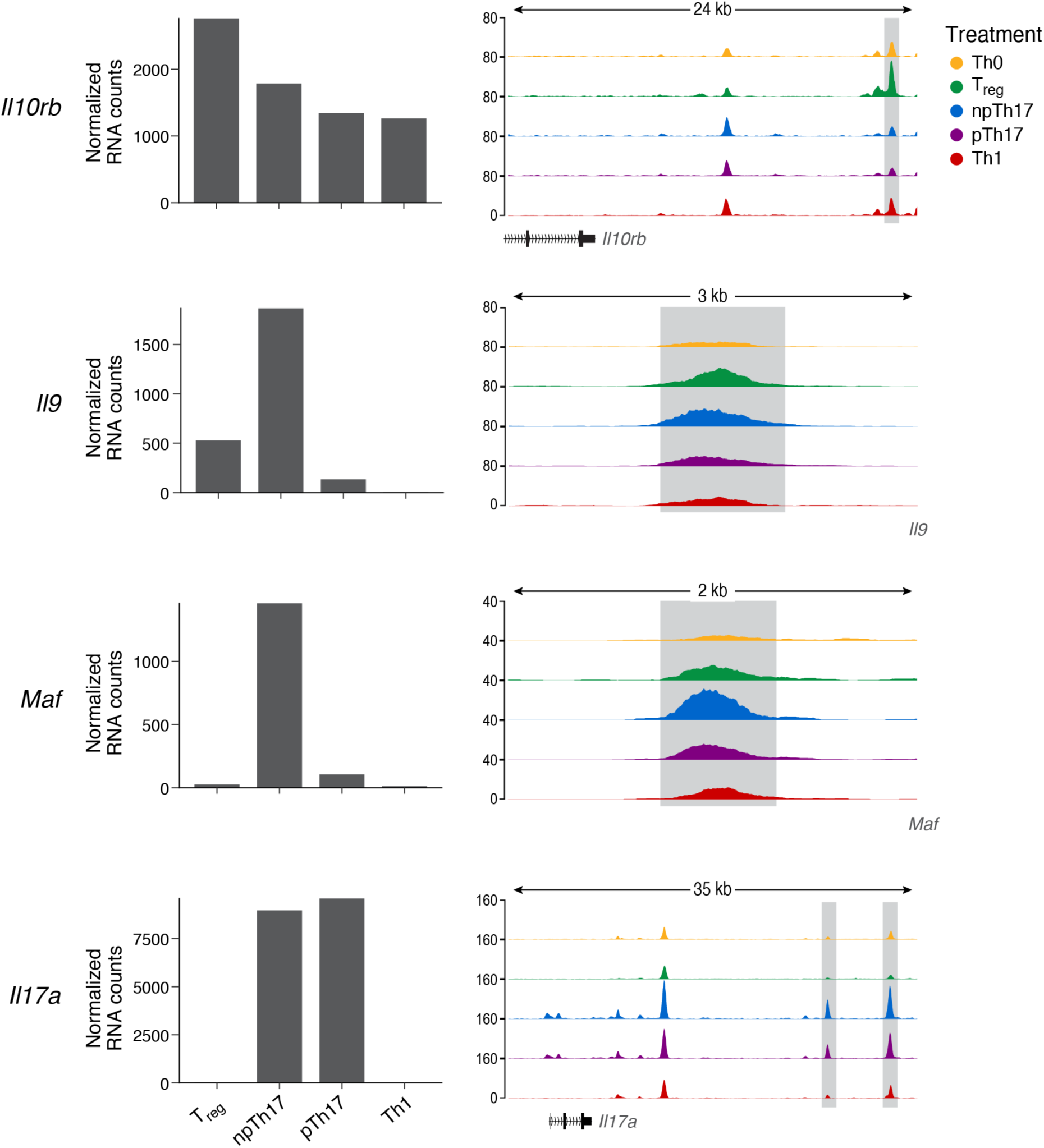
ATAC-seq peaks across CD4^+^ T cell subsets. RNA expression level (left, *y* axis, normalized transcript counts) and enrichment of ATAC-seq signal over background (right, *y* axis, log fold) of *Il10rb* (chr16:91,420,666-91,445,000), *Il9* (chr13:56,505,989-56,508,498), *Maf* (chr8:115,567,448-115,569,757), *Il17a* (chr1:20,727,418-20,762,267) loci in in each CD4^+^ T cell subset. Gene bodies are displayed on the bottom. In cases of distal peaks the associated gene is indicated (right: downstream).

**Fig. S5:**
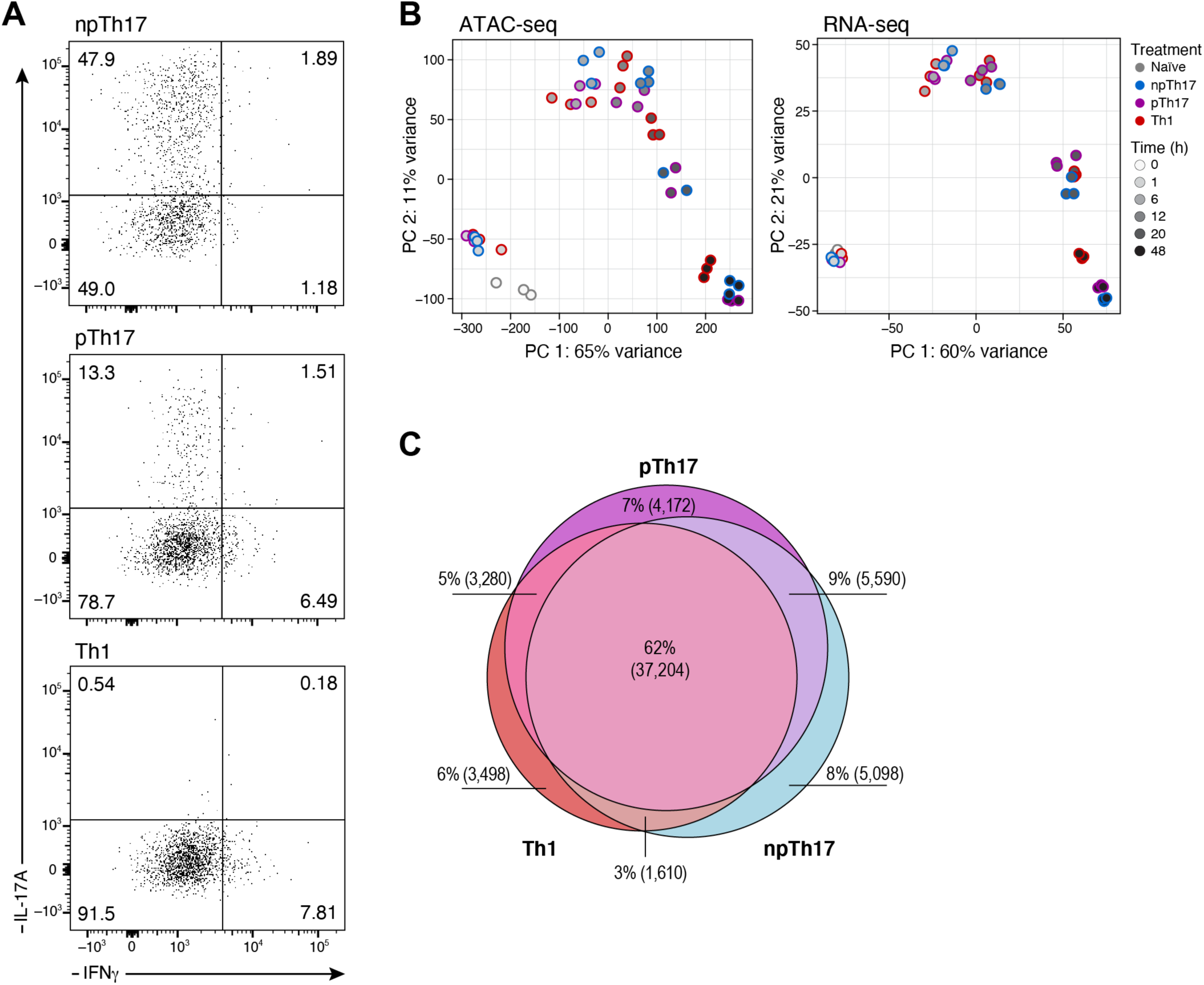
Time course analysis of chromatin accessibility during Th17 and Th1 differentiation. (**A**) Expression of the marker cytokines at 48h. Flow cytometry plots for expression of IL-17A (*y* axis) and IFN𝛾 (*x* axis) at 48h during the time course experiment. (**B**) Time and activation are captured by independent axes of variation (PCs) in both bulk ATAC- and RNA-seq. PCA plot of PC1 and PC2 of bulk ATAC-seq (left) and bulk RNA-seq (right) of naïve and *in vitro* differentiated npTh17 (blue), pTh17 (purple) and Th1 (red) cells at 0h, 1h, 6h, 12h, 20h, and 48h (grey scale). (**C**) Shared chromatin regions close during Th17 and Th1 differentiation. Venn diagram of chromatin regions that close by 48h in npTh17 (blue), pTh17 (purple) or Th1 (red) cells.

**Fig. S6:**
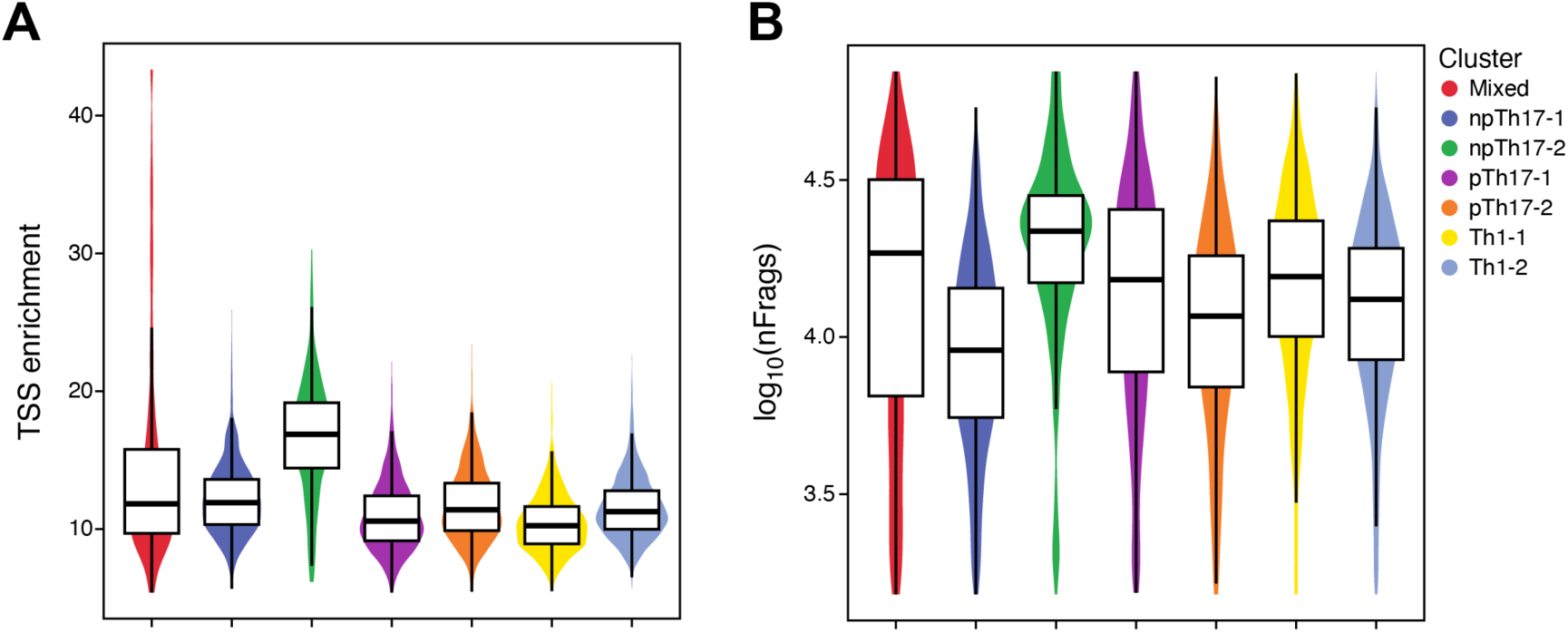
Quality control metrics of scATAC-seq. Distribution of TSS enrichment (**A,** *y* axis) and number of fragments (**B**, *y* axis) for each scATAC-seq cluster (*x* axis) (as in Fig. 5B).

**Fig. S7:**
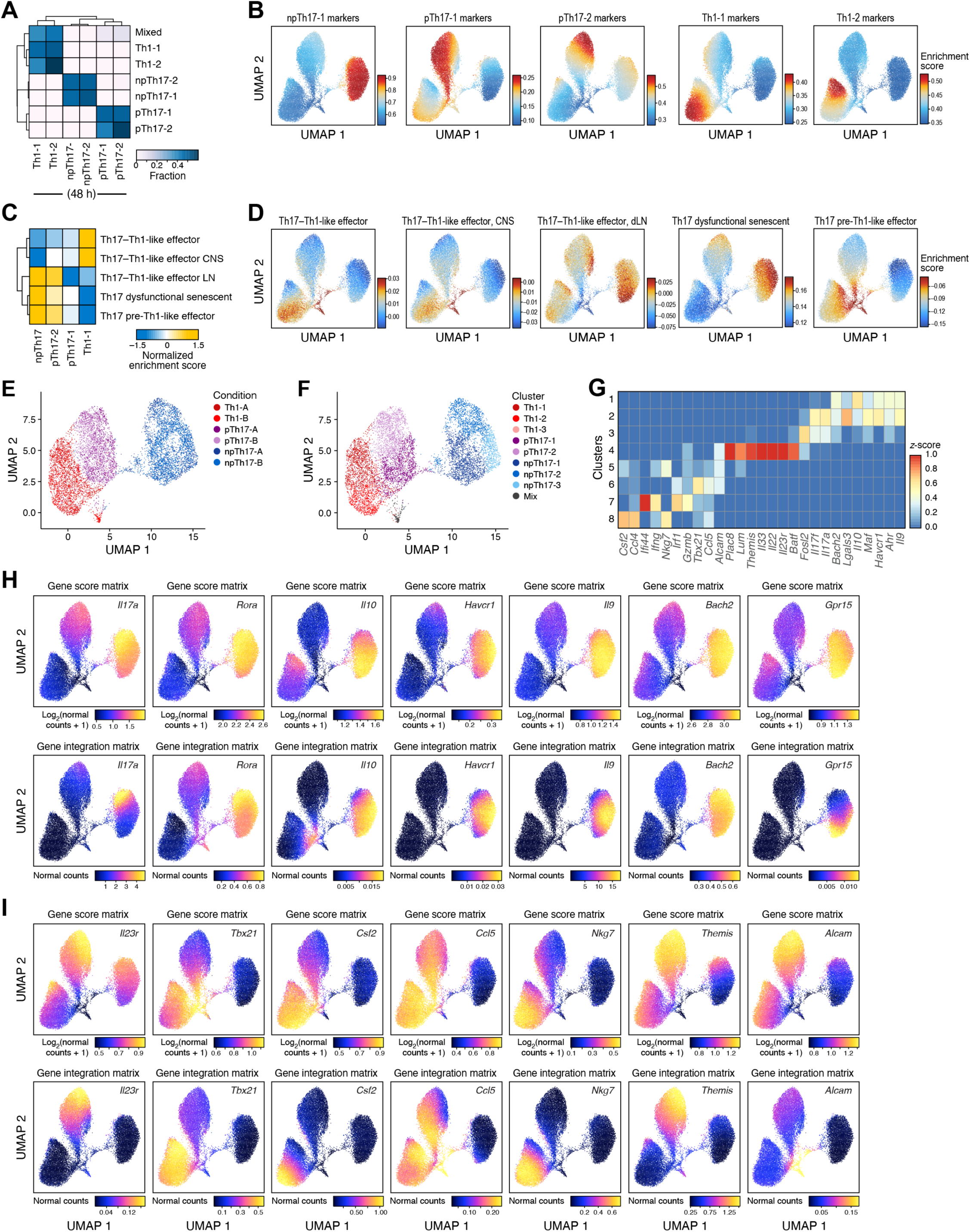
ScATAC-seq and scRNA-seq of Th17 cells. (**A,B**) ScATAC-seq profiles cluster by cell subset. (**A**) Fraction of cells (color bar) of each sample (columns) that are members of each scATAC-seq cluster (rows, defined as in Fig. 5B). (**B**) UMAP embedding of scATAC-seq profiles (as in Fig. 5B) colored by enrichment score of cluster markers (as in Fig. 5C). (**C,D**) ScATAC-seq profile clusters are enriched for distinct Th17 cell signatures. (**C**) Enrichment (color bar) of Th17 pathway signatures ^8^ (rows) with genes associated with differentially accessible ATAC peaks in each scATAC-seq cluster (columns). (**D**) UMAP embedding of scATAC-seq profiles (as in Fig. 5B) colored by enrichment score of Th17 pathway signatures. (**E,F**) ScRNA-seq of *in vitro* differentiated npTh17, pTh17, and Th1 cells at 48h. UMAP embedding of scRNA-seq profiles colored by treatment (**E**) or cluster assignment (**F**). (**G**) Differential peak-to-gene association of key Th17 effector genes in distinct scATAC-seq clusters. Relative signal (z-score, color bar) of peak-to-gene links for selected Th17 effector genes (columns) in each cluster (rows). (**H,I**) Variation in chromatin accessibility of npTh17 and pTh17 marker genes across scATAC-seq profiles. UMAP embedding of scATAC-seq profiles (as in Fig. 5B) colored by ATAC-signal as gene score (top) and predicted gene expression as gene integration (bottom) for selected marker genes of npTh17 (**H**) and pTh17 (**I**) cells.

**Fig. S8:**
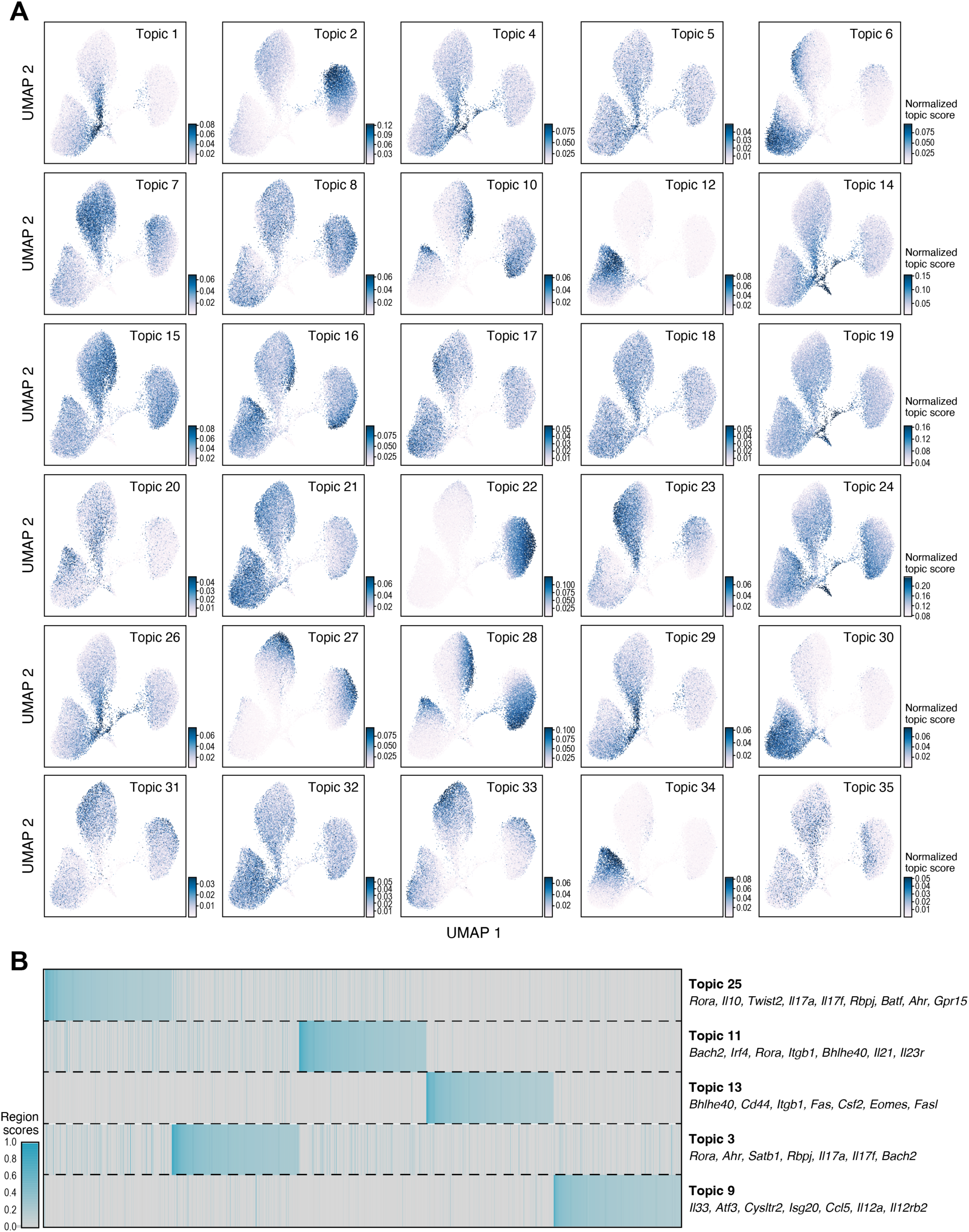
ScATAC-seq topic modeling of Th17 and Th1 cells. (**A**) Distinct and shared gene programs across Th17 and Th1 cells identified by topic modeling. UMAP embedding of scATAC-seq profiles (dots) colored by topic scores (normalized topic score, colorbar) for 30 of 35 topics (the remaining 5 topics are in Fig. 5F). (**B**) Chromatin peaks underlying key topics. Regions scores (colorbar) for each of the top 500 accessible chromatin peaks (columns) in five selected topics (rows, same topics as in Fig. 5F). Selected genes associated with peaks in each topic are listed on the right.

**Fig. S9:**
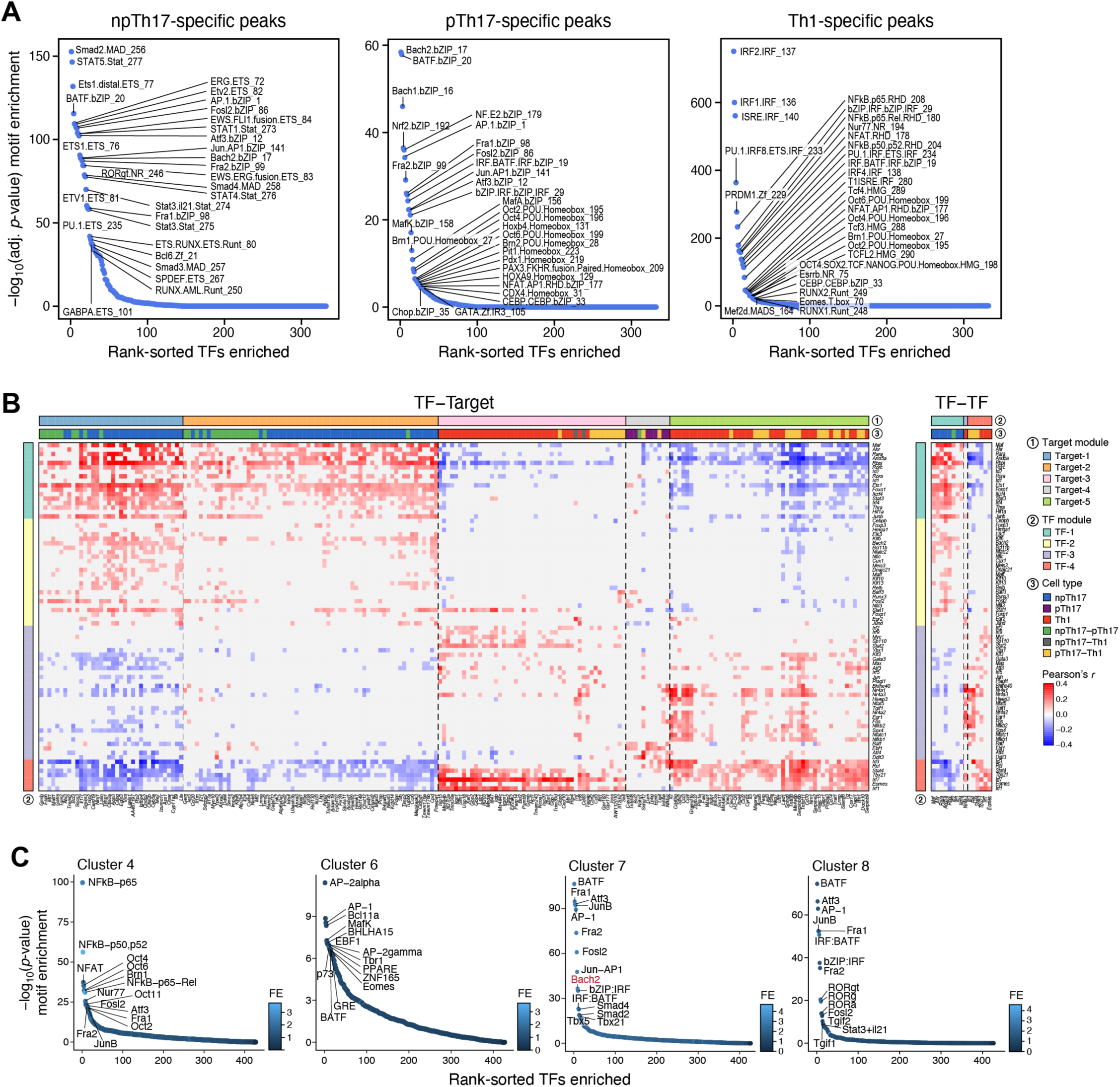
TF motif enrichment in scATAC-seq peaks. (**A**) TF motifs enriched in accessible peaks in different cell subsets. Significance (- log_10_(Bonferoni adjusted p-value), *y* axis) and rank (*x* axis) of enrichment of motifs (dots) in npTh17- (left), pTh17- (middle), and Th1-specific (right) peaks. (**B**) Comprehensive inferred Th17 and Th1 regulatory network. Pearson correlation coefficients (red/blue color bar) between each TF in the network (row) and each target (column) for target genes that are (right) or are not (left) themselves TFs in the network. TFs and targets are clustered (color bars 1 and 2) and the cell type(s) where an interaction is detected are labeled (color bar 3). (**C**) Transcription factor motif enrichment of DACR clusters. Significance (-log_10_(Bonferroni adjusted p-value), *y* axis), rank (*x* axis), and fold enrichment compared to background peak set (dot color) of enrichment of motifs (dots) in DACR clusters.

**Fig. S10:**
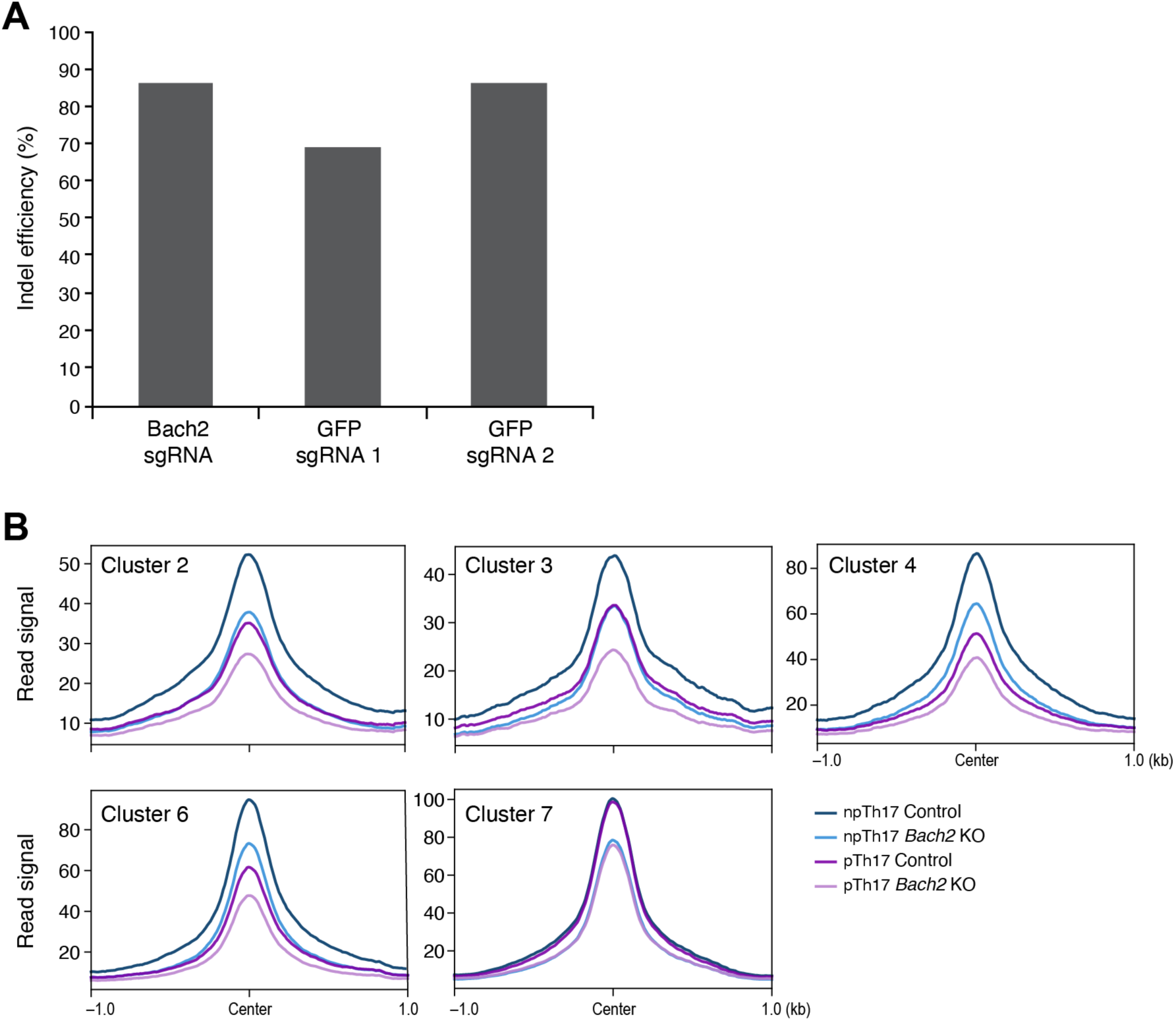
Validation of *Bach2* CRISPR/Cas9 editing efficiency. (**A**) Editing efficiency at the *Bach2* locus. Editing efficiency (*y* axis; % loci with indels by Sanger sequencing) for a Bach2-targeting sgRNA (measured at the *Bach2* locus) and for two sgRNAs targeting the GFP transgene locus (*x* axis). (**B**) Impact of *Bach2* KO on chromatin accessibility in T_reg_ and Th17 related loci. Global chromatin accessibility (*y* axis, average normalized reads) in npTh17 (blue) and pTh17 (purple) cell populations from *Bach2* KO (light color) or control (dark color) cells in chromatin regions that are more accessible in T_reg_-specific clusters (2-4 in Fig. 3E) and in Th17-specific clusters (6,7 in Fig. 3E).

**Fig. S11:**
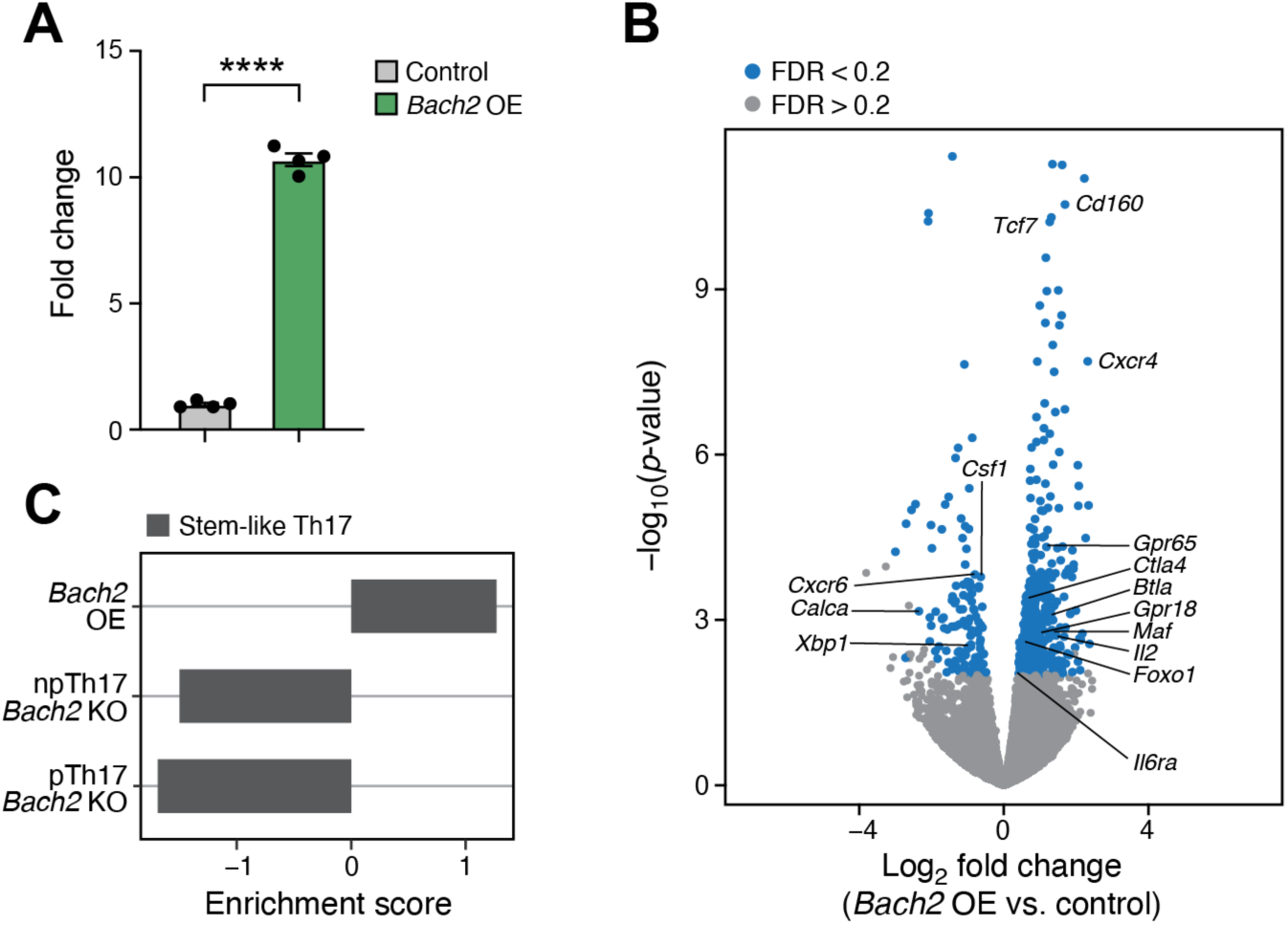
B*a*ch2 overexpression in Th17 cells. (**A**) *Bach2* overexpression validation. *Bach2* transcript expression (*y* axis, fold change by qPCR, mean ± SEM) in *Bach2* overexpressing (*Bach2* OE) and control pTh17 cells (*x* axis). ****, P <0.0001, unpaired two-tailed t-test. (**B**) Differentially expressed genes in *Bach2* OE pTh17 2D2 cells. Significance (-log_10_(p-value), *y* axis) and effect size (log_2_(fold change), *x* axis) of differential expression of each gene between transduced cells (GFP^+^) isolated from the CNS of recipients of *Bach2* OE or control pTh17 2D2 cells. Blue: FDR <0.2. (**C**) BACH2 induces a stem-like Th17 cell program. Enrichment score (*x* axis) of the *in vivo* stem-like Th17 cell signature ^16^ with genes associated with differentially accessible ATAC peaks in the *Bach2* OE and *Bach2* KO cells (*y* axis).

**Fig. S12:**
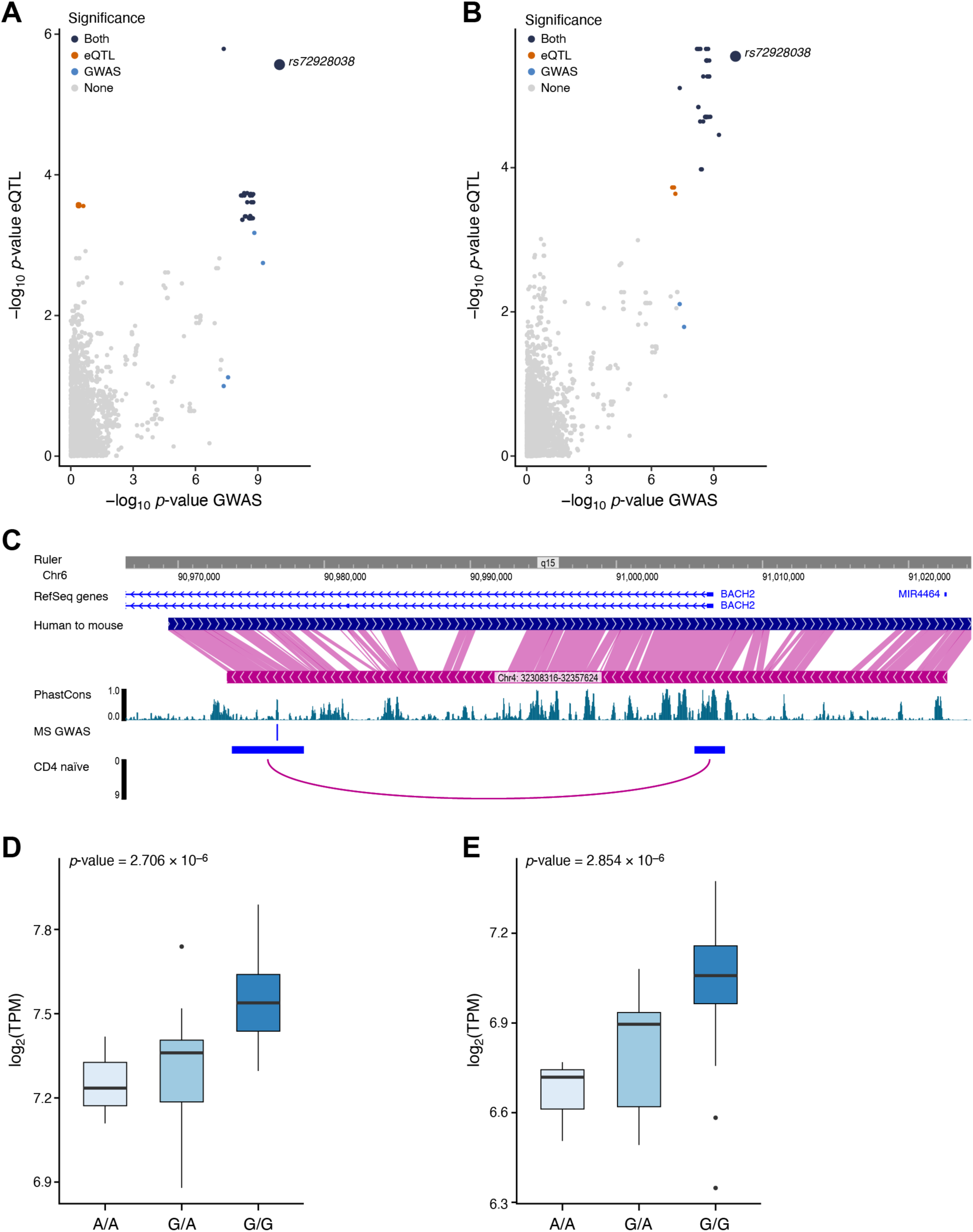
Association of MS-associated variant rs72928038 with increased BACH2 expression. (**A,B**) Rs72928038 is a variant associated with MS by GWAS and a BACH2 eQTL in T cells. Significance (–log10(p-value)) of association of each variant (dot) with MS risk in GWAS (*x* axis; from the discovery phase of IMSGC ^77^) and as a BACH2 cis-eQTL (*y* axis, from the database of immune cell expression study [DICE] ^80^) in CD4^+^ naïve (**A**) or CD4^+^ T_reg_ (**B**) cells for variants in a region spanning 1MB up and downstream of rs72928038. Rs72928038 post-replication p = 8.38*10^-29^ and odds ratio for G allele was 0.866. Variants are colored by statistical significance in both MS GWAS and eQTL study (dark blue), only in MS GWAS (light blue), only in eQTL study (light red), or neither (light grey). Statistical significance thresholds: MS GWAS: P <5*10^-8^ (genome wide); eQTL: FDR <10% (locus only test). (**C**) Putative regulatory relationship of rs72928038 with BACH2 expression. Tracks from top to bottom display: position in chromosome 6; reference sequence genes (truncated view); alignment of mouse genome (mm9) to human genome (hg19; https://vizhub.wustl.edu/public/hg19/weaver/hg19_mm9_axt.gz); evolutionary conservation in 46 vertebrate species; position of rs72928038; and promoter capture interaction loop between enhancer intronic region, overlapping rs72928038, and BACH2 promoter in CD4^+^ naïve cells ^79^. All positions are in human genome version 19. (**D,E**) Association of rs72928038 variant with increased BACH2 expression. Distribution of gene expression (*y* axis, log_2_(transcripts per million) (log_2_(TPM)) for each rs72928038 genotype (*x* axis) in CD4^+^ naïve cells (**D**) and CD4^+^ T_reg_ cells (**E**). P-values are from the respective cis-eQTL analysis of rs72928038 and BACH2 ^80^.

